# A genetically encoded redox bottleneck constrains human developmental rate

**DOI:** 10.64898/2026.09.10.750701

**Authors:** Gabriel E. Valdebenito, Jack P. Madden, Carolina Mendez, Leslie M. Azurdia, Xintao Zhu, Siyu Chen, Diana Atlas, Katherine Introcaso, Hao Zhang, Yahui Wang, Fangcong Dong, Rohit Sharma, Vamsi K. Mootha, Gary J. Patti, Margarete Diaz-Cuadros

## Abstract

The intrinsically slow pace of human development poses challenges for regenerative medicine and disease modeling. This trait is attributed to low metabolic rates, yet the endogenous mechanisms determining species-specific metabolic flux remain unknown. Here, we identify coupling between glycolytic NADH production and mitochondrial oxidation through the glycerol-3-phosphate (G3P) shuttle as a genetic bottleneck constraining human developmental tempo. Using stem cell-derived models of the segmentation clock, an oscillator whose period reflects developmental rate, we show that low expression of the G3P shuttle enzyme GPD1L limits NADH oxidation in human progenitors compared to mouse. Overexpressing GPD1L boosts metabolic flux, accelerating the segmentation clock, cell cycle, and differentiation across germ layers. G3P-mediated redox coupling is thus a genetically encoded, rate-limiting mechanism that sets the tempo of human development.

## Main Text

Among mammals, the rate of embryonic development is widely divergent(*1, 2*). Despite following the same sequence of developmental events, human embryos progress through embryogenesis at roughly half the rate as mouse embryos(*3*). These tempo differences are intrinsic to cells and are recapitulated *in vitro*(*4–7*). For instance, human and mouse stem cell-derived models display distinct oscillation periods of the segmentation clock(*4, 6*). This molecular oscillator governs the rhythmic formation of somites in the presomitic mesoderm (PSM)(*8–13*) and provides a tractable system to study species-specific developmental rate.

Interspecies variation in basal metabolic rates influence developmental tempo(*14–24*), with elevated metabolic rates giving rise to accelerated development. This regulation is potentially mediated by nicotinamide adenine dinucleotide (NAD⁺/NADH) redox metabolism, and exogenously altering redox balance can modulate developmental rate *in vitro*(*21*). Redox metabolism plays a fundamental role in regulating key cellular processes, including energy production, signal transduction, and the maintenance of redox homeostasis(*25, 26*). At the core of these functions lies the NAD⁺/NADH redox couple, which acts not only as an essential cofactor in metabolic reactions but also as a critical regulator of metabolic flux(*24, 27*). The intracellular NAD⁺/NADH ratio influences several metabolic nodes: glycolytic enzymes converting glucose into pyruvate, the pyruvate dehydrogenase complex enabling mitochondrial entry, and enzymes in the TCA cycle and electron transport chain that drive oxidative phosphorylation(*28–30*). Here we set out to identify the endogenous mechanisms that set species-specific metabolic rates and NAD⁺/NADH ratios, which ultimately determine developmental rate.

## Results

### NAD^+^/NADH ratio regulates developmental rate

To explore whether redox metabolism regulates developmental rate, we utilized pluripotent stem cell (PSC)-derived models of the mouse and human segmentation clock(*4, 21*). Under identical culture conditions, both species expressed comparable primed pluripotency markers (**fig. S1, A-D**) and robustly differentiated into presomitic mesoderm (PSM) (**fig. S1E**), upregulating expected lineage markers including *MSGN1*, *TBXT*, *TBX6* (**fig. S1, F and G**), and *HOX* genes (**fig. S1H**)(*4, 21, 31–33*). Consistent with previous reports, mouse cells displayed significantly faster developmental kinetics than human cells, characterized by earlier *Msgn1* reporter induction (24 h vs 48 h, **fig. S2, A-D**)(*4, 34*) and a two-fold faster segmentation clock period (2.7 h vs 5.0 h, **Fig. 1A, fig. S2, E-H**)(*2, 4–6, 21, 35–37*).

**Fig. 1.**
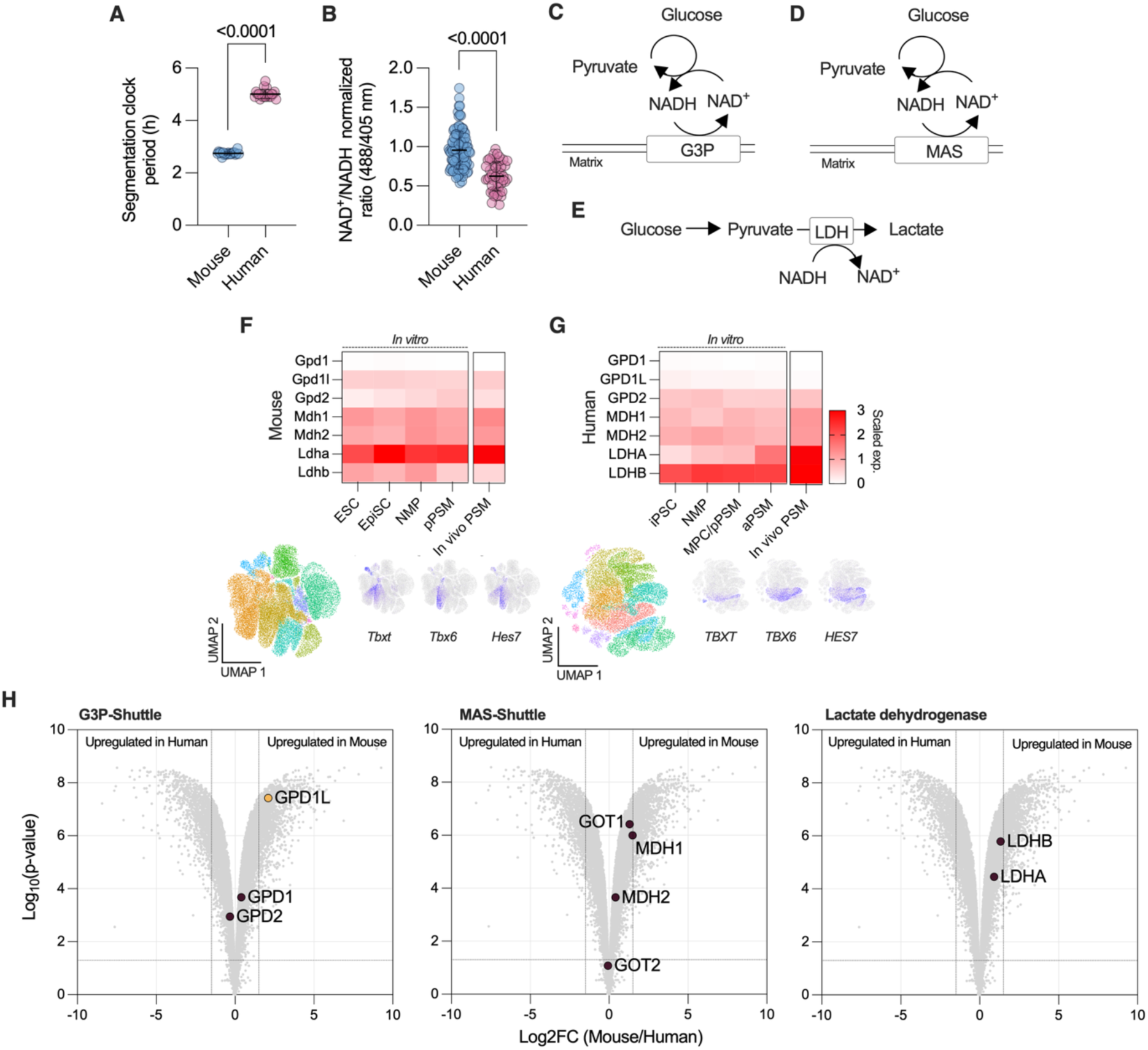
Divergence in NAD^+^ regeneration pathways between mouse and human presomitic mesoderm. (**A**) Segmentation clock period in pluripotent stem cell-derived mouse and human PSM cells as measured by *Hes7-Achilles* fluorescent reporter, establishing a two-fold difference in oscillatory pace (20 data points from three independent replicates; mean ± s.d.; unpaired two-tailed t-test, *P* < 0.0001). (**B**) Ratiometric imaging of SoNar excitation (488/405nm) as a proxy for NAD⁺/NADH levels in mouse and human pluripotent stem cell-derived PSM cells, indicating a more oxidized redox balance in mouse cells (n > 3 independent replicates; > 200 cells per replicate; mean ± s.d.; unpaired two-tailed t-test, *P* < 0.0001). (C–E) Schematic representation of the three major cytosolic NADH-oxidation routes: the glycerol-3-phosphate (G3P) shuttle (**C**), the malate–aspartate shuttle (MAS) (**D**), and lactate dehydrogenase (LDH)-mediated conversion (**E**). Matrix refers to the mitochondrial matrix. (F–G) Top: Heatmaps showing scaled expression of genes encoding components of the three NADH-oxidation routes in single cell RNA-sequencing data from mouse (**F**) and human (**G**) pluripotent stem cells differentiating towards PSM fate, or *in vivo* PSM cells from mouse and human embryos (data from 3 independent replicates per species)(*4, 50, 51*). NMP = neuromesodermal progenitors, pPSM = posterior PSM, MPC = mesodermal progenitor cells, aPSM = anterior PSM. Bottom: UMAPs of single cell RNA-sequencing data for mouse E8.0 (**F**) and human PCW3 (**G**) embryos, colored by cluster. Heatmaps on the right show the expression of PSM marker genes. (**H**) Volcano plots comparing the bulk transcriptomes of mouse and human *in vitro*-derived PSM for genes in each of the three NADH-oxidation pathways. The cytosolic G3P dehydrogenase *GPD1L* shows reduced expression in human PSM cells relative to mouse (data from 3 independent replicates per species; statistical analysis as specified in Methods). Vertical and horizontal lines indicate significance thresholds (log₂ fold change > 1.5 and adjusted P < 0.05).

Our previous work implicated the cellular redox state in regulating developmental tempo across species(*21*). We previously observed that the extracellular [pyruvate]/[lactate] ratio, an indirect indicator of the cytosolic NAD⁺/NADH balance, was higher in mouse than in human PSM cells(*21*). To directly quantify the cytosolic redox state, we expressed and validated the genetically encoded NAD⁺/NADH sensor SoNar(*38, 39*) **(fig. S3, A and B)**, which shows differential excitation profiles when bound to NAD^+^ or NADH. SoNar quantification confirmed that mouse PSM cells maintain a significantly higher cytosolic NAD⁺/NADH ratio than human cells **(Fig. 1B and fig. S3C)**.

We next tested whether direct modulation of the NAD^+^/NADH ratio by genetic means could alter the developmental rate of human PSM cells. We expressed a water-forming bacterial oxidase, *Lb*NOX(*40*), and a soluble transhydrogenase from *Escherichia coli*, *Ec*STH(*41, 42*), to manipulate the cytosolic NAD⁺/NADH ratio in human cells. *Lb*NOX uses NADH as a substrate to produce NAD⁺ and H₂O, whereas *Ec*STH interconverts NAD⁺ and NADPH to NADH and NADP⁺ **(fig. S3D).** As previously reported(*21*), *Lb*NOX expression increased the NAD⁺/NADH ratio relative to control cells **(fig. S3E)**, which was accompanied by accelerated PSM cell fate acquisition as measured by MSGN1-YFP induction **(fig. S3, F and H)** and a shortened segmentation clock oscillation period by ∼30 minutes **(fig. S3, I and J**). By contrast, *Ec*STH expression decreased the cytosolic NAD⁺/NADH ratio **(fig. S3E)**, decelerated MSGN1-YFP induction **(fig. S3, G and H)**, and lengthened the segmentation clock oscillation period by ∼20 minutes **(fig. S3, I and K)**. Taken together, these data established that direct modulation of the cytosolic NAD⁺/NADH ratio influences both differentiation kinetics and segmentation clock periodicity in human PSM cells.

### G3P-shuttle uncoupling in human PSM cells

We next set out to define the molecular basis underlying the distinct redox state of mouse and human PSM cells. In cells, the cytosolic NAD⁺/NADH ratio is maintained by three major systems that transfer reducing equivalents to the mitochondria or regenerate NAD⁺ locally: the glycerol-3-phosphate (G3P) shuttle, the malate–aspartate (MAS) shuttle, and lactate dehydrogenase (LDH) **(Fig. 1C-E)**(*43–45*). The G3P shuttle oxidizes cytosolic NADH through GPD1 or GPD1L, which reduces dihydroxyacetone phosphate (DHAP) to G3P(*44, 46*). This metabolite then transfers electrons to mitochondrial FAD through GPD2, regenerating NAD⁺ in the cytosol while feeding electrons to the electron-transport chain **(Fig. 1C)**. In parallel, the MAS shuttle mediates a malate– aspartate cycle in which MDH1 and MDH2 interconvert malate and oxaloacetate across the mitochondrial compartments, while GOT1 and GOT2 catalyze the corresponding transamination steps between aspartate and glutamate(*47*), together providing another route for cytosolic NADH oxidation **(Fig. 1D)**. Finally, the LDH reaction reduces pyruvate to lactate using NADH, directly regenerating NAD⁺ in the cytoplasm without transferring reducing power to mitochondria(*48*). Although this pathway efficiently restores redox balance, it does so at the expense of carbon utilization, as lactate is exported from the cell, leading to a net loss of carbon available for biosynthetic metabolism(*44, 48, 49*) **(Fig. 1E)**.

To compare the expression dynamics of these enzymes across developmental stages, we re-analyzed our published single-cell RNA-sequencing dataset(*4*) spanning mouse and human pluripotent stem cells, neuromesodermal progenitors, and presomitic mesoderm *in vitro* **(Fig. 1, F and G)**. MAS- and LDH-associated genes (*Mdh1/2*, *Ldha/b*) were robustly expressed in both species. In mouse, both the cytosolic *Gpd1l* and the mitochondrial *Gpd2* components of the G3P shuttle were expressed **(Fig. 1F)**. In contrast, human cells expressed mitochondrial *GPD2* but displayed minimal or undetectable levels of the cytosolic G3P-shuttle enzyme *GPD1L* **(Fig. 1G).** Neither species expressed measurable levels of *GPD1* (**Fig. 1, F and G**). This pattern suggested a species-specific uncoupling of cytosolic NADH oxidation from mitochondrial electron transfer in human cells. We corroborated these findings by analyzing *in vivo* single-cell RNA-sequencing datasets from E8.0 (embryonic day 8.0) mouse(*50*) and PCW3 (post-conception week 3) human embryos(*51*). Clusters expressing PSM markers (*TBXT, TBX6, HES7*) confirmed that the species-specific G3P-shuttle expression pattern observed *in vitro* reflects real differences between mouse and human embryos **(Fig. 1, F and G; lower panels)**.

We next sought to confirm these observations in our *in vitro* differentiated cells using bulk RNA-sequencing. Replicates clustered tightly within each species and showed marked separation between mouse and human samples (**fig. S4, A and B**), with differential expression analysis of 1-to-1 orthologs identifying thousands of transcripts displaying significant interspecies variation (**fig. S4C**). Analysis of NADH-oxidation genes showed that while MAS- and LDH-related genes displayed similar expression across species, the cytosolic G3P-shuttle component *GPD1L* was strongly upregulated in mouse cells (**Fig. 1H**). These results suggest that the G3P shuttle is active in mouse PSM cells as both the cytosolic (*Gpd1l*) and mitochondrial (*Gpd2*) components are present, thereby forming a coupled shuttle that can run as a cycle. In contrast, human PSM cells express the mitochondrial enzyme GPD2 but largely lack the cytosolic component (*GPD1* or *GPD1L*), meaning that the shuttle is uncoupled and likely not functional. Mouse PSM cells may therefore maintain an active cytosolic–mitochondrial NADH shuttle via G3P, while human PSM cells lack this coupling, potentially contributing to the lower NAD⁺/NADH ratio and slower developmental tempo observed in the human lineage.

To assess whether the transcriptional differences in G3P-shuttle gene expression translated into functional divergence, we first confirmed differential protein expression of the corresponding enzymes **(Fig. 2A)** and validated these patterns in PSM cells derived from one additional mouse and two additional human embryonic stem cell lines **(fig. S5A)**. We next performed targeted metabolomic analysis in cells cultured for 24 h in either unlabeled or uniformly labeled [U-¹³C_6_]glucose to quantify steady-state metabolite abundance and isotopic enrichment. Overall, metabolite abundance and labeling patterns across glycolysis, the TCA cycle, lactic fermentation and glutaminolysis were broadly comparable between species **(fig. S5B)**.

**Fig. 2.**
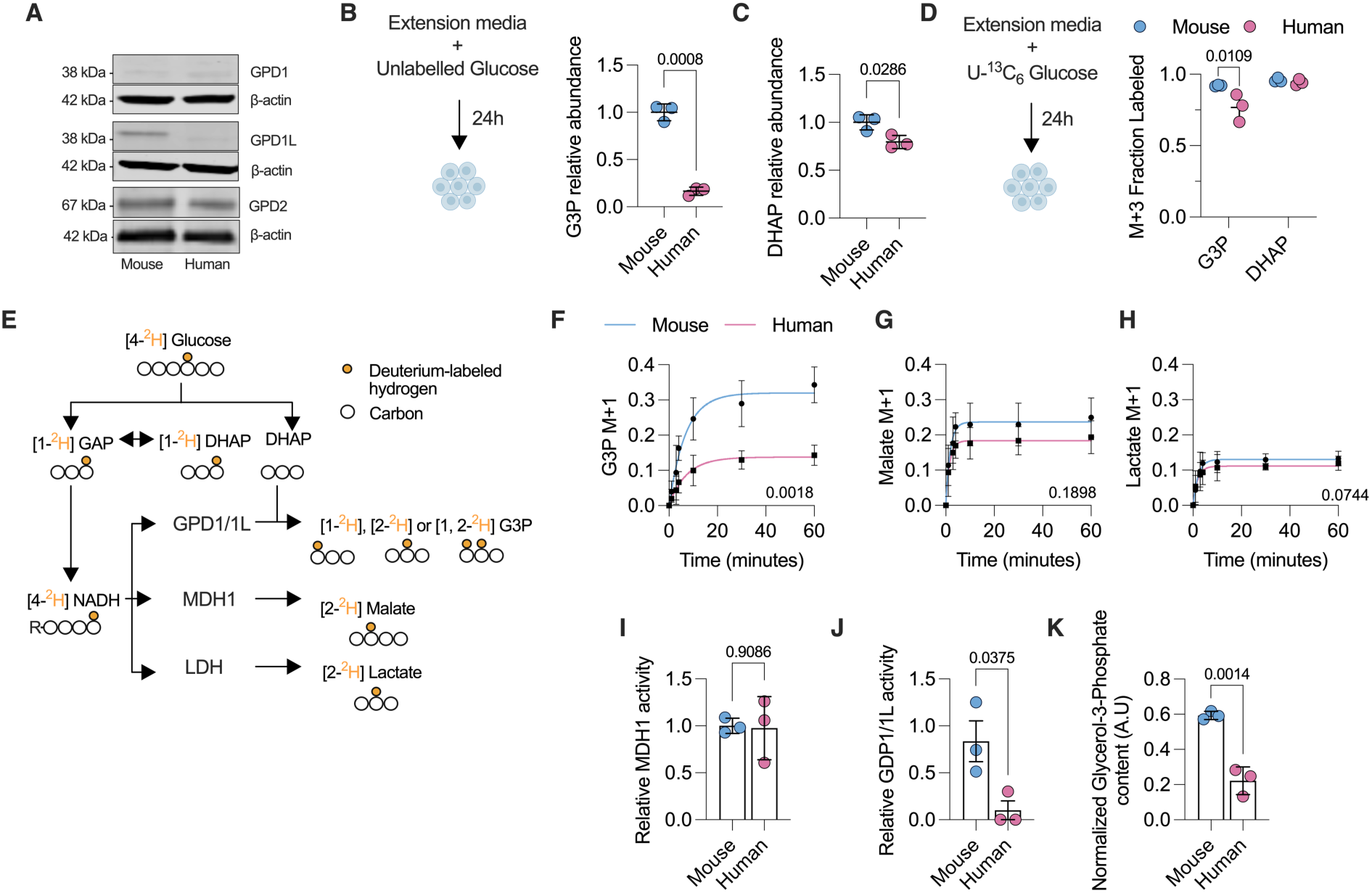
The Glycerol-3-Phosphate shuttle is functionally uncoupled in human PSM cells. (**A**) Immunoblot analysis showing cytosolic (GPD1, GPD1L) and mitochondrial (GPD2) glycerol-3-phosphate dehydrogenases in mouse and human pluripotent stem cell-derived PSM cells (n = 3 independent replicates). (**B**) Experimental schematic (left) illustrating the steady-state metabolite analysis after 24-hour incubation with unlabeled glucose, used to quantify glycerol-3-phosphate (G3P) abundance (right) in mouse and human *in vitro*-derived PSM cells (n = 3 independent replicates; mean ± s.d.; unpaired two-tailed t-test, *P* = 0.0008). (**C**) Relative abundance of dihydroxyacetone phosphate (DHAP) in mouse and human *in vitro*-derived PSM cells after 24 h incubation with unlabeled glucose, showing a modest decrease in human cells (n = 3 independent replicates; mean ± s.d.; unpaired two-tailed t-test, *P* = 0.0286). (**D**) Schematic of the [U-¹³C_6_]glucose tracing experiment (left) and corresponding M+3 isotopologue fractions of G3P and DHAP (right) in mouse and human *in vitro*-derived PSM cells. Human cells exhibit reduced M+3 labeling in G3P, while DHAP labeling remains comparable between species, consistent with attenuated G3P-shuttle activity (mean ± s.d.; n = 3 independent experiments per species; unpaired two-tailed t-test; G3P, *P* = 0.0109; DHAP, ns). (**E**) Diagram of the [4-²H]glucose tracing strategy. During glycolysis, the deuterium label is transferred to NADH at the GAPDH step and subsequently appears in NADH-dependent reduction products (G3P, malate, lactate). (**F–H**) Time-course of deuterium enrichment (M+1) in G3P (**F**), malate (**G**), and lactate (**H**) following [4-²H]glucose labeling at different timepoints for mouse and human *in vitro*-derived PSM cells. Mouse cells show markedly faster G3P labeling kinetics, whereas malate and lactate labeling exhibit minimal interspecies differences (mean ± s.e.m.; n = 3 independent replicates; repeated-measures two-way ANOVA with Tukey correction; G3P, *P* = 0.0018; malate, ns; lactate, ns). (**I–J**), Relative enzymatic activities of MDH1 (**I**) and GPD1/GPD1L (**J**) inferred from the initial slopes of G3P (**F**) and malate (**G**) labeling curves (n = 3 independent replicates; mean ± s.d.; unpaired two-tailed t-test; i, ns; j, *P* = 0.0375). (**K**) Quantification of normalized glycerol-3-phosphate (G3P) content as measured by enzymatic assay in mouse and human pluripotent stem cell-derived PSM cells, showing reduced steady-state levels in human PSM cells (n = 3 independent replicates; mean ± s.d.; unpaired two-tailed t-test;lis *P* = 0.0014).

Comprehensive metabolite profiling further showed broadly similar abundance across amino-acid, carbohydrate, and lipid metabolism, whereas nucleotide-associated metabolites and redox cofactors were generally higher in mouse cells, and amino-acid metabolites tended to be higher in human cells **(fig. S5C)**. Among all metabolites measured, those directly linked to cytosolic redox reactions showed the strongest interspecies differences: human cells exhibited a pronounced depletion of G3P, the product of the cytosolic G3P-shuttle reaction, consistent with their low GPD1/GPD1L expression **(Fig. 2B)**. In contrast, the upstream metabolite DHAP showed only a modest decrease **(Fig. 2C)**, suggesting that the deficit reflects reduced conversion rather than substrate limitation.

^1^³C label incorporation analysis revealed comparable M+3 enrichment in DHAP between species, indicating that glucose catabolism through upper glycolysis is largely conserved **(Fig. 2D)**. In contrast, G3P showed slightly lower M+3 labeling in human cells, consistent with reduced conversion of DHAP to G3P through cytosolic G3P-shuttle activity **(Fig. 2D)**. These findings show that the loss of G3P in human cells reflects a specific impairment in NADH-dependent reduction rather than a generalized reduction in glycolytic throughput.

Whereas uniformly labeled [U-¹³C] glucose tracing captures the flow of carbon through glycolysis, it does not directly report on redox transfer. To specifically quantify cytosolic NADH oxidation, we therefore performed an orthogonal stable-isotope tracing experiment using [4-²H] glucose(*43*). In this assay, a deuterium atom from the C-4 position of glucose is transferred to NADH by glyceraldehyde 3-phosphate dehydrogenase (GAPDH). The labeled hydrogen can subsequently appear in downstream reduction products formed by NADH-dependent reactions, providing a direct readout of cytosolic redox turnover **(Fig. 2E).** Upon NADH oxidation in the cytoplasm, the deuterium label is transferred to either G3P (via GPD1/GPD1L), malate (via MDH1 in the MAS shuttle), or lactate (via LDH). Whereas malate M+1 and lactate M+1 did not show differences in labeled fractions over time between species, G3P showed significantly higher labeling in mouse cells than human PSM cells **(Fig. 2F-H).** These results suggest that the G3P shuttle is significantly less active in human PSM cells, consistent with transcriptomic and immunoblotting data.

We next used deuterium labeling patterns to estimate relative enzymatic activities in each species(*43*). This revealed comparable MDH1 activity (**Fig. 2I)** but reduced GPD1/GPD1L activity in human cells (**Fig. 2J)**. To independently validate these findings biochemically, we quantified intracellular glycerol-3-phosphate levels, which were markedly lower in human cells (**Fig. 2K**), consistent with diminished G3P-shuttle function.

Altogether, these results strongly suggest that species-specific enzyme expression patterns give rise to elevated G3P shuttle activity in mouse PSM cells relative to human cells. The G3P shuttle may therefore underlie the observed divergent redox balance between mouse and human PSM cells, linking cytosolic NADH recycling to differences in developmental tempo.

### Elevated glycolytic flux in mouse PSM cells

Having identified the G3P shuttle as a key regulator of cytosolic redox state, we hypothesized that the absence of this shuttle in human cells might impose a bottleneck on metabolic throughput. Like cancer cells, embryonic cells rely on aerobic glycolysis (i.e. the Warburg effect) to sustain high rates of growth and proliferation(*52–54*). During glycolysis, NAD^+^ is reduced to NADH by GAPDH **(fig. S6A)**. Continued glycolytic flux requires the oxidation of NADH back to NAD^+^(*48, 55, 56*). We reasoned that a more reduced cytosolic redox balance may limit glycolytic flux in human PSM cells. Indeed, previous studies found that human PSM cells displayed lower levels of glycolytic activity compared to mouse cells(*21, 35, 57*). To corroborate these findings with a direct readout of glycolytic function, we performed a glycolytic stress test using the Seahorse extracellular flux analyzer **(fig. S6B)**. Upon glucose addition, mouse cells exhibited a more rapid and robust increase in extracellular acidification rate (ECAR), indicative of higher basal glycolytic activity compared to human cells **(fig. S6, B and C)**. Subsequent oligomycin treatment further elevated ECAR in both cell types, reflecting enhanced glycolytic capacity under mitochondrial ATP synthase inhibition. However, the increase was markedly greater in mouse cells, indicating higher glycolytic reserve and capacity (**fig. S6, B and C)**. The final addition of 2-deoxy-D-glucose (2DG) suppressed ECAR to baseline levels, confirming that the acidification was primarily due to glycolysis. These results suggest that mouse cells possess a greater glycolytic capacity than human PSM. To independently verify enhanced glycolytic output, we quantified extracellular lactate accumulation over time. Mouse PSM cultures secreted significantly more lactate than human cells **(fig. S6D)**, consistent with their higher glycolytic flux measured by ECAR. Altogether, these data led us to conclude that mouse PSM cells experience higher glycolytic flux than human PSM cells.

We next performed a comparative transcriptomic analysis of mouse and human PSM cells to investigate whether species-specific gene expression patterns reflect the observed differences in glycolytic flux and redox balance. Gene Ontology (GO) enrichment analysis revealed strong over-representation of glycolytic processes in mouse cells, including *glucose catabolic process to pyruvate*, *canonical glycolysis* and *NADH regeneration* **(fig. S6E)**. Expression profiling of glycolytic genes further revealed selective upregulation of a subset of genes in mouse PSM relative to human cells **(fig. S6F)**, consistent with elevated glycolytic flux. We also observed species-specific gene expression differences in pathways controlling NADH utilization and NAD⁺ regeneration, including NADH-linked tricarboxylic-acid (TCA)-cycle enzymes, pyruvate-dehydrogenase complex activity, amino-acid and branched-chain metabolism, NAD⁺ biosynthesis, and NAD⁺-consuming reactions **(fig. S7A)**. We additionally examined curated metabolic pathway gene sets and performed KEGG pathway enrichment analysis **(fig. S7, B and C)**. Both approaches highlighted a consistent enrichment of glycolysis, NADH regeneration, oxidative phosphorylation, and TCA-cycle pathways in mouse cells, whereas human PSM showed relative enrichment of signaling, translational, and biosynthetic processes. Species-specific transcriptomic profiles therefore support our observation that mouse PSM cells display a more glycolytic phenotype compared with human PSM.

### Shuttle manipulation alters PSM tempo

Interspecies differences in gene expression, metabolite abundance, and metabolic flux prompted us to test whether introducing mouse-like metabolic features into human cells could accelerate cell fate acquisition and segmentation clock dynamics. To this end, we overexpressed ***GPD1*** or ***GPD1L*** in human PSM cells. Transcriptional activation of the endogenous human *GPD1 or GPD1L* loci was achieved using a CRISPR activation (CRISPRa) system in which sgRNAs recruited a dCas9–VP64 complex to activate target gene expression **(Fig. 3A)**(*58–60*). Immunoblotting and tandem mass tag (TMT)-based proteomics confirmed robust overexpression of GPD1 and GPD1L **(fig. S8, A and B).**

**Fig. 3.**
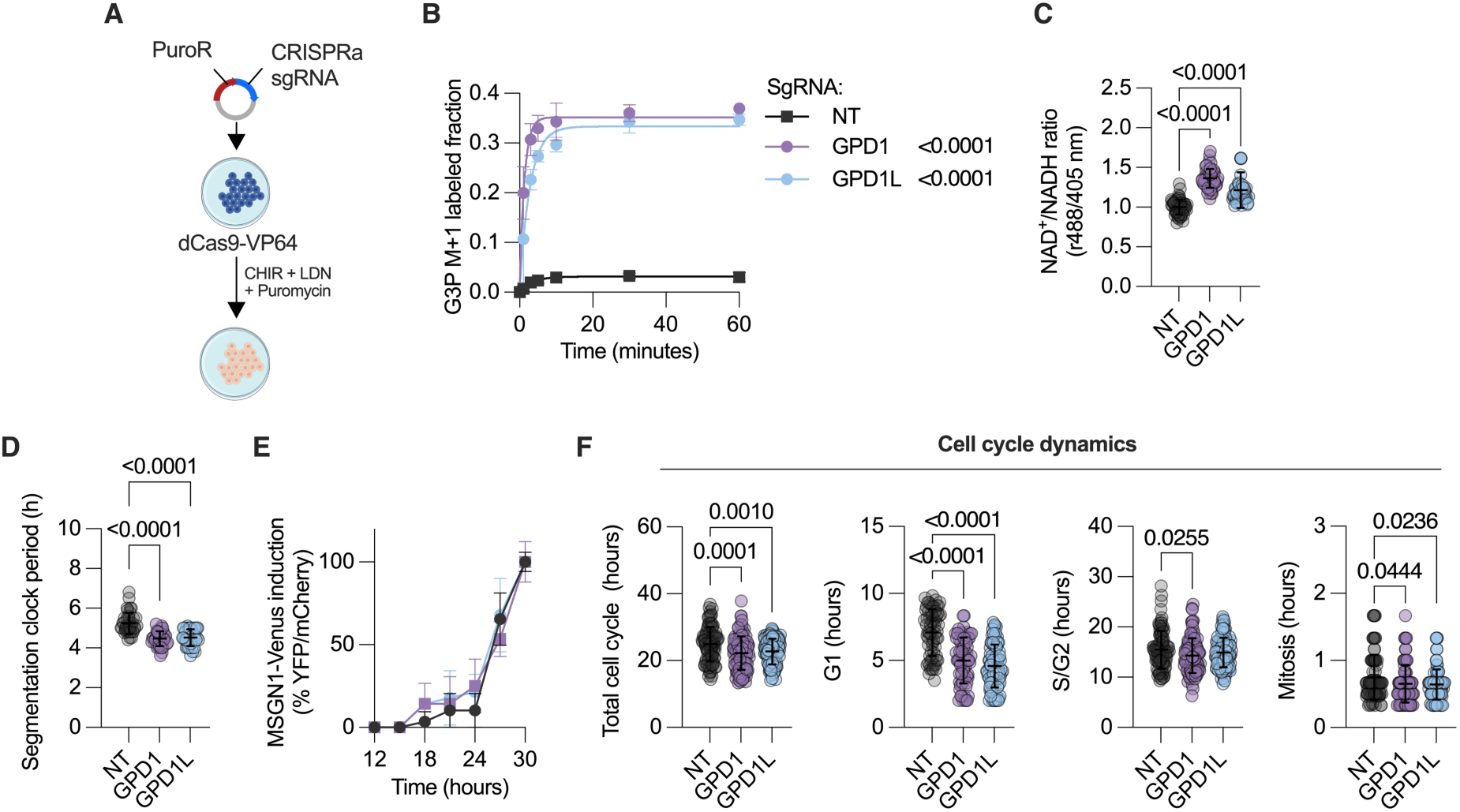
Activation of GPD1 or GPD1L enhances NADH oxidation and accelerates developmental tempo in human PSM. (**A**) Schematic of CRISPRa strategy using dCas9-VP64 and sgRNAs targeting *GPD1* or *GPD1L*. (**B**) Time-course of deuterium enrichment (M+1) in glycerol-3-phosphate (G3P) following [4-²H]glucose labeling in human PSM cells expressing sgRNAs targeting *GPD1* or *GPD1L* for CRISPRa overexpression, showing increased labeling kinetics upon *GPD1* or *GPD1L* activation relative to a non-targeting (NT) sgRNA control (n = 3 independent replicates; repeated-measures two-way ANOVA with Tukey correction; GPD1/NT, *P* < 0.0001; GPD1L/NT, *P* < 0.0001). (**C**) Quantification of cytosolic NAD⁺/NADH ratio by SoNar ratiometric imaging in control (NT) and *GPD1/GPD1L*-overexpressing human PSM cells *in vitro*, demonstrating enhanced NADH oxidation upon *GPD1*- and *GPD1L activation* (n > 3 independent replicates; > 200 cells per replicate; mean ± s.d.; one-way ANOVA with Tukey correction; GPD1/NT, *P* < 0.0001; GPD1L/NT, *P* < 0.0001). (**D**) Segmentation clock period measured using the *HES7–Achilles* reporter in control (NT) and *GPD1/GPD1L*-overexpressing human PSM cells *in vitro,* revealing accelerated oscillations in *GPD1*- and *GPD1L*-activated cells relative to non-targeting (NT) controls (n > 30 oscillation traces from 3 independent experiments; mean ± s.d.; one-way ANOVA with Tukey correction; GPD1/NT, P < 0.0001; GPD1L/NT, P < 0.0001). (**E**) Representative traces of *MSGN1–Venus* induction dynamics in differentiating human iPSCs *in vitro* under control (NT) or *GPD1/GPD1L-*overexpression conditions (n = 3 independent replicates; repeated-measures two-way ANOVA with Tukey correction; GPD1/NT, ns; GPD1L/NT, ns). (**F**) FUCCI cell-cycle profiling in control (NT) and *GPD1/GPD1L*-overexpressing human PSM cells *in vitro,* showing shortened total and G₁-phase durations with modest reductions in S/G₂ and mitosis upon *GPD1* or *GPD1L* activation compared to non-targeting (NT) controls (n > 50 cells per replicate; 3 independent replicates; mean ± s.d.; one-way ANOVA with Tukey correction; total cell cycle, GPD1/NT, *P* = 0.0001, GPD1L/NT, *P* = 0.0010; G₁ phase, GPD1/NT, *P* < 0.0001, GPD1L/NT, *P* < 0.0001; S/G₂ phase, GPD1/NT, *P* = 0.0255, GPD1L/NT, ns; mitosis, GPD1/NT, *P* = 0.0444, GPD1L/NT, *P* = 0.0236).

We next assessed whether this overexpression was sufficient to functionally activate the G3P shuttle using the aforementioned deuterium tracing strategy. GPD1 and GPD1L (*GPD1/1L)*-overexpressing cells displayed a rapid increase in the M+1 G3P isotopologue compared to non-targeting controls **(Fig. 3B)** without changes in malate or lactate labeling **(fig. S8, C and D)**, indicating selective activation of the G3P shuttle.

Ratiometric SoNar imaging showed an elevated NAD⁺/NADH ratio in overexpressing cells **(Fig. 3C)**, confirming enhanced cytosolic NADH oxidation. This redox shift coincided with faster segmentation clock oscillations, shortening the period by approximately 40–50 minutes **(Fig. 3D and fig. S8, E and F)**. Despite this acceleration, the timing of MSGN1-YFP induction remained unchanged in GPD1/1L overexpressing cells **(fig. 3E)**. This may reflect the dual signaling and bioenergetic role of glycolytic flux in mesoderm specification(*61*). Because the metabolic state often influences proliferation(*62*), we examined whether G3P-shuttle activation affected the cell cycle. Using FUCCI reporter expressing cells **(fig. S8G)**, we observed a pronounced shortening of the G1 phase and a corresponding reduction in total cycle length **(Fig. 3F**), demonstrating that enhanced redox flux and metabolic activity accelerate both oscillatory and proliferative tempos.

We next evaluated whether the increased NADH oxidation capacity imparted by GPD1/1L overexpression led to increased glycolytic flux in human PSM cells. Indeed, we observed higher levels of basal and maximal glycolytic capacity in cells overexpressing GPD1 and GPD1L compared to control **(Fig. 4, A and B and fig. S8H)**. Because the G3P shuttle links cytosolic NAD^+^ regeneration with electron transport chain activity, we also evaluated mitochondrial metabolism to determine whether G3P-shuttle activation impacts cellular respiration. *GPD1/1L*-overexpressing cells exhibited significantly higher basal and maximal oxygen-consumption rates **(Fig. 4, C and D and fig. S8I)**, consistent with elevated mitochondrial utilization of glycolysis-derived NADH and pyruvate. Importantly, we confirmed that GPD1/1L overexpression gave rise to increased complex I-independent respiration **(Fig. 4E)**. This reflects the activity of the mitochondrial arm of the G3P shuttle, whereby GPD2 donates electrons directly to ubiquinone through FADH_2_. Recoupling of the G3P shuttle in human PSM cells therefore simultaneously boosts both glycolytic flux and mitochondrial respiration.

**Fig. 4.**
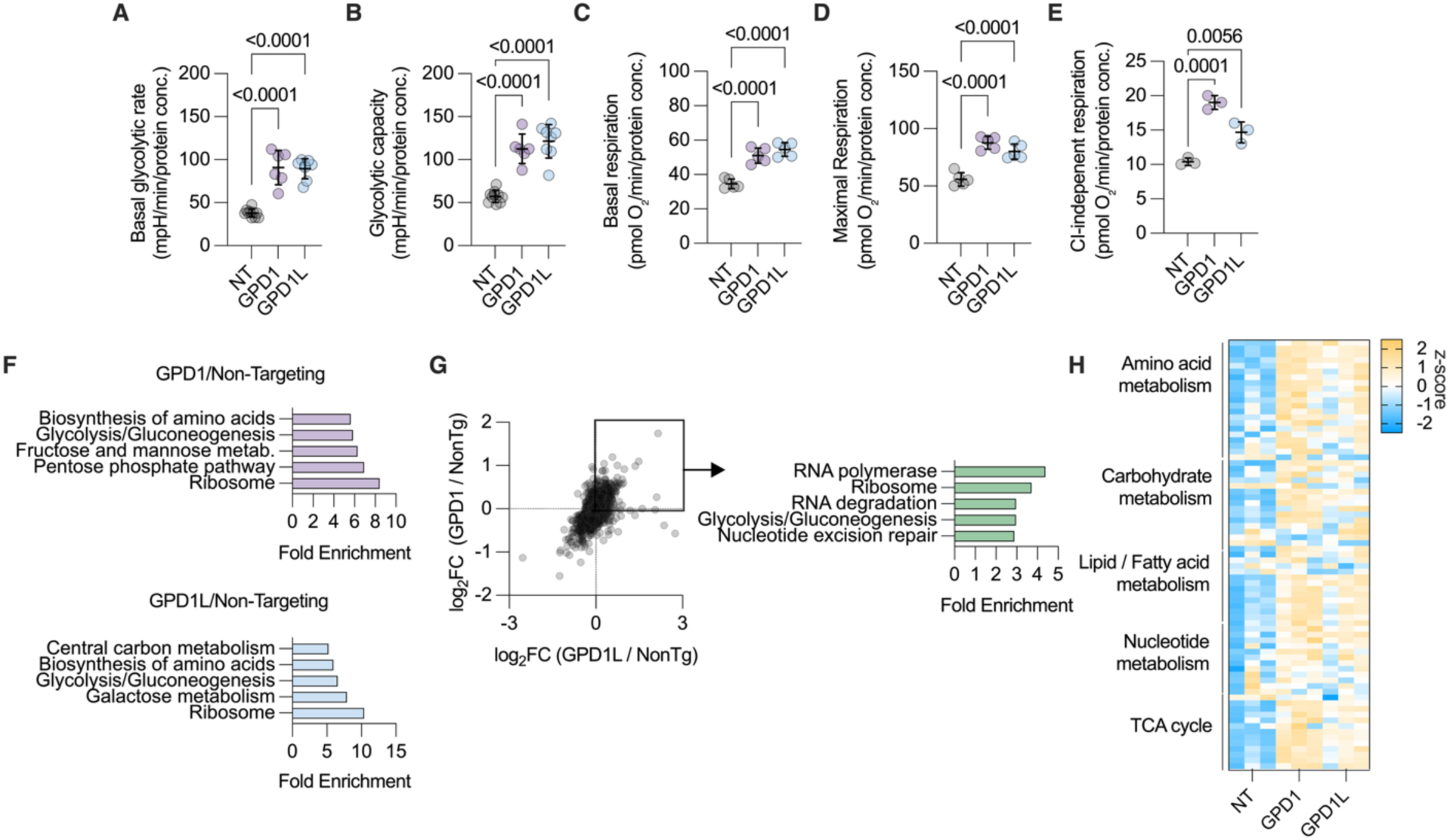
G3P shuttle activation is sufficient to accelerate glycolysis and mitochondrial respiration in human PSM. (**A–B**) Basal glycolytic rate (**A**) and glycolytic capacity (**B**) in control (NT) and GPD1/GPD1L-overexpressing human PSM cells *in vitro* (n = 3 independent replicates; mean ± s.d.; one-way ANOVA with Tukey correction; basal glycolytic rate, GPD1/NT, *P* < 0.0001, GPD1L/NT, *P* < 0.0001; glycolytic capacity, GPD1/NT, *P* < 0.0001, GPD1L/NT, *P* < 0.0001). (**C–D**) Oxygen-consumption rate (OCR) analysis in control (NT) and *GPD1/GPD1L*-overexpressing human PSM cells *in vitro,* showing increased basal (**C**) and maximal (**D**) respiration upon *GPD1* or *GPD1L* activation compared to non-targeting (NT) controls (n = 6 independent replicates; mean ± s.d.; one-way ANOVA with Tukey correction; basal respiration, GPD1/NT, *P* < 0.0001, GPD1L/NT, *P* < 0.0001; maximal respiration, GPD1/NT, *P* < 0.0001, GPD1L/NT, *P* < 0.0001). (**E**) Complex I (CI)–independent respiration in control (NT) and *GPD1/GPD1L*-overexpressing human PSM cells *in vitro,* showing elevated non-NADH-linked oxygen consumption in *GPD1*- and *GPD1L*-activated cells (n = 3 biologically independent samples per condition; mean ± s.d.; one-way ANOVA with Tukey correction; GPD1/NT, *P* = 0.0001; GPD1L/NT, *P* = 0.0056). (**F**) Gene Ontology (GO) enrichment of transcripts up-regulated in human PSM cells *in vitro* by *GPD1* or *GPD1L* activation relative to non-targeting controls, highlighting glycolytic, biosynthetic, and ribosomal pathways. (**G**) Scatter plot (left) comparing log₂-fold changes between GPD1 and GPD1L activation in human PSM cells based on proteomics analysis, showing concordant up-regulation of glycolytic and NADH-regeneration pathways by gene ontology analysis (right). (**H**) Heat map showing metabolite abundance (z-score) in human PSM cells *in vitro* across non-targeting (NT), *GPD1*, and *GPD1L* CRISPRa conditions (n = 3 independent replicates per condition).

To better understand the changes in cellular physiology caused by G3P shuttle activation, we conducted transcriptomic, proteomic, and targeted metabolomic profiling of GPD1/1L-overexpressing cells. We then performed pathway-enrichment analysis comparing GPD1- and GPD1L-overexpressing cells to non-targeting controls. Both conditions were enriched for glycolysis/gluconeogenesis, amino-acid biosynthesis, and ribosome pathways **(Fig. 4F)**, indicating coordinated upregulation of anabolic and energy-producing processes. *GPD1* overexpression produced stronger enrichment of pentose-phosphate and fructose/mannose metabolism, whereas *GPD1L* preferentially affected central-carbon and galactose metabolism, suggesting partially overlapping but distinct metabolic rewiring. To determine the shared proteomic consequences of *GPD1* and *GPD1L* activation, we compared fold-changes between both conditions **(Fig. 4G)**. Proteins upregulated in both contexts were enriched for glycolytic enzymes, ribosomal components, and RNA-processing factors, suggesting that G3P-shuttle activation enhances metabolic and biosynthetic capacity at the protein level. Grouping of detected metabolites by major biochemical pathways revealed coordinated increases across amino-acid, carbohydrate, lipid/fatty-acid, nucleotide, and TCA-cycle metabolism **(Fig. 4H)**, consistent with a global enhancement of central-carbon metabolism and biosynthetic output.

### Three-lineage acceleration by redox modulation

Because GPD1/1L overexpression accelerated the segmentation clock within the mesodermal lineage, we next asked whether our observations also extended to other germ layers. To direct pluripotent stem cells toward the definitive endoderm (DE) lineage, Nodal/Activin and WNT signaling pathways were activated using Activin A and CHIR99021, followed by immunostaining for SOX17, a hallmark of endodermal specification **(Fig. 5A)**. To generate neural progenitor cells (NPCs), Nodal and BMP signaling were inhibited using dual-SMAD inhibition with SB431542 and LDN193189, resulting in the induction of PAX6 expression, a canonical marker of neuroectodermal identity **(Fig. 5B)**. In both contexts, mouse cells differentiated more rapidly than human cells, consistent with lineage-independent differences in developmental tempo.

**Figure 5.**
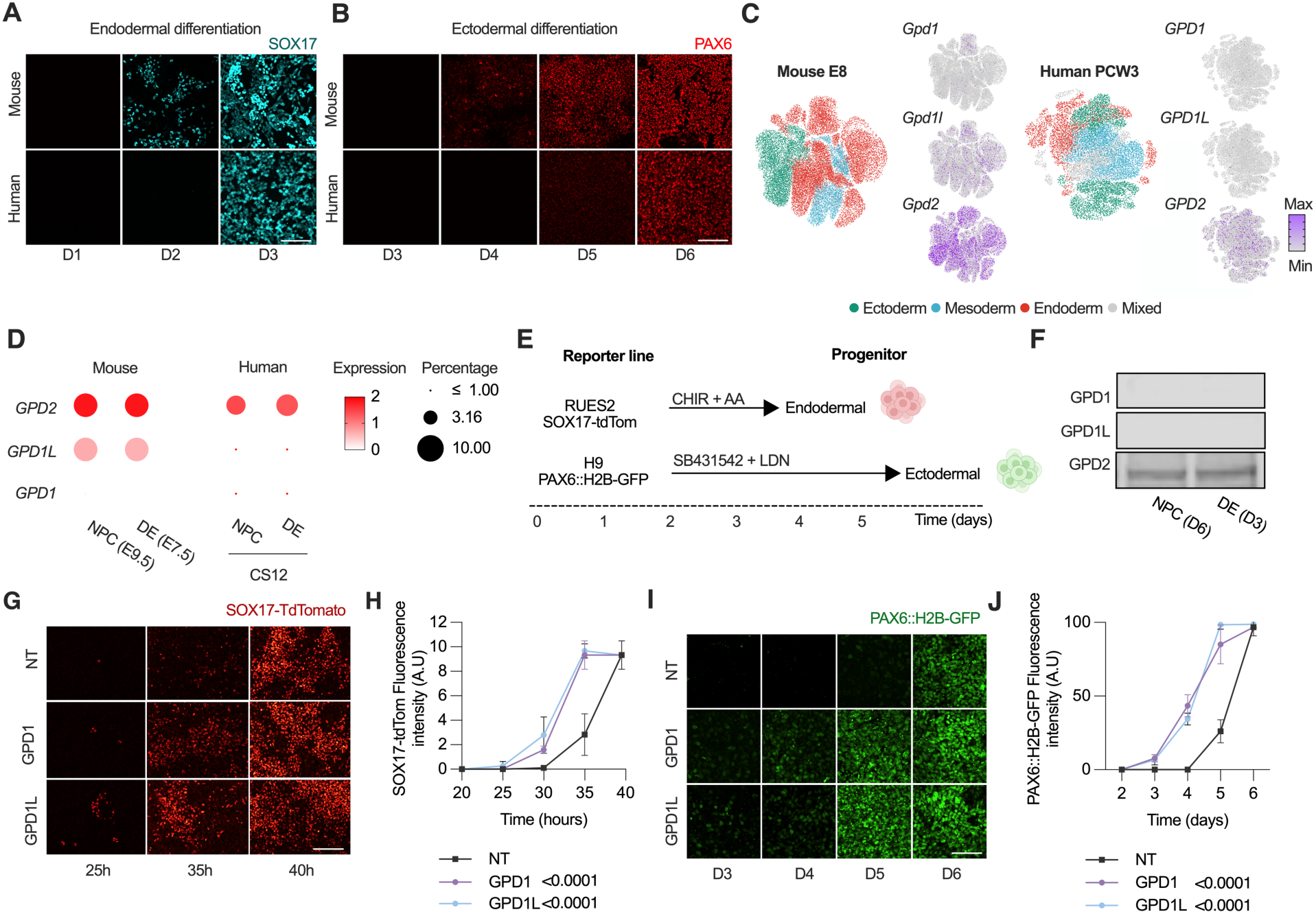
Species-specific tempo differences and redox modulation extend across germ-layer lineages. (**A**) Representative images of SOX17 immunofluorescence during definitive endoderm differentiation of mouse and human pluripotent stem cells, showing earlier induction in mouse cultures. D1-D3 = Day1-Day3. Scale bar, 100 µm. (**B**) Representative images of PAX6 immunofluorescence during ectodermal differentiation of mouse and human pluripotent stem cells, revealing faster neuroectodermal specification in mouse relative to human cultures. Scale bar, 100 µm. (**C**) UMAP of single-cell RNA-seq data from E8.0 mouse and PCW3 human embryos showing expression of *Gpd1*, *Gpd1l*, and *Gpd2 (right)* across ectodermal, mesodermal, and endodermal derivatives (color coded on the left)(*50, 51*). (**D**) Dot-plot showing expression of *GPD1*, *GPD1L* and *GPD2* in mouse (E7.5–E9.5) and human (Carnegie stage S12) single-cell transcriptomes corresponding to clusters of neural progenitor cell (NPC) and definitive endoderm (DE) identities. Dot size represents the percentage of cells expressing each gene, and color intensity indicates average expression level (normalized log-scale). Data from published single-cell RNA-seq atlases of early embryogenesis. (**E**), Schematic of germ-layer differentiation protocols and human embryonic stem cell reporter lines used: *RUES2 SOX17-tdTomato* (endoderm) and *H9 PAX6-H2B-GFP* (ectoderm). (**F**) Immunoblot analysis showing expression of GPD1, GPD1L, and GPD2 across human lineages *in vitro*: definitive endoderm (DE, Day 3 of differentiation), and neural progenitor cells (NPC, Day 6 of differentiation). (**G**) Representative fluorescence images of SOX17–tdTomato reporter signal during definitive endoderm differentiation of human pluripotent stem cells overexpressing GPD1 or GPD1L compared with non-targeting (NT) controls, showing accelerated SOX17 induction in GPD1/GPD1L cells. Time points indicate hours after differentiation onset. Scale bar, 100 µm. (**H**) Quantification of SOX17–tdTomato fluorescence intensity over time (n = 3 independent replicates; mean ± s.d.; one-way ANOVA with Tukey correction; GPD1/NT, *P* < 0.0001; GPD1L/NT, *P* < 0.0001). (**I**) Representative fluorescence images of PAX6::H2B–GFP reporter signal during neural progenitor differentiation of human pluripotent stem cells overexpressing GPD1 or GPD1L compared with non-targeting (NT) controls, showing earlier onset of PAX6 expression in GPD1/GPD1L cultures. Time points indicate days after induction. Scale bar, 100 µm. (**J**) Quantification of PAX6::H2B–GFP fluorescence intensity over time (n = 3 independent replicates; mean ± s.d.; one-way ANOVA with Tukey correction; GPD1/NT, *P* < 0.0001; GPD1L/NT, *P* < 0.0001).

To evaluate whether the species-specific G3P shuttle gene expression pattern detected in PSM applies to other germ layers, we reanalyzed single-cell RNA-seq datasets from E8.0 mouse and PCW3 human embryos, clustering cells by germ-layer identity. The G3P-shuttle expression phenotype observed in PSM extended across all germ layers: mouse cells exhibited higher expression of *Gpd1l* and *Gpd2* compared with their human counterparts **(Fig. 5C),** with little *Gpd1* expression in either species. To connect these *in vivo* observations with *in vitro* differentiation, we next selected NPC and DE populations based on marker gene expression. Both cell types displayed the expected species-specific pattern of G3P shuttle gene expression (**Fig. 5D)**.

We next tested whether enhancing cytosolic NADH oxidation within these two cell fates could accelerate differentiation speed. To this end, we used PAX6::GFP(*63*) and SOX17-tdTomato(*64*) human reporter lines to monitor neuroectodermal and endodermal induction, respectively **(Fig. 5E)**. These reporter lines did not express GPD1 or GPD1L at the protein level at day 6 and day 3 of ectodermal and endodermal induction, respectively **(Fig. 5F)**, demonstrating that *in vitro* differentiation recapitulates the phenotype observed in PSM cells. The reporter lines were then engineered to stably express dCas9–VP64 and subjected to CRISPRa-mediated activation of GPD1 or GPD1L, with overexpression verified by immunostaining **(fig. S9)**. Strikingly, GPD1/GPD1L overexpression led to earlier activation of lineage reporters: SOX17-tdTomato-positive cells were first detected one day earlier, and PAX6::GFP-positive cells approximately two days earlier, relative to non-targeting controls **(Fig. 5G-J)**.

Together, these findings suggest that activation of the G3P shuttle broadly accelerates human differentiation kinetics across germ layers, indicating that metabolic redox coupling serves as a general determinant of developmental tempo rather than a lineage-restricted regulator.

## Discussion

We identified GPD1L as a metabolic regulator of species-specific developmental tempo, revealing a direct link between cytosolic redox state and the pace of embryogenesis. Using a unified human– mouse pluripotent stem cell platform with identical culture conditions, we eliminated extrinsic variables and uncovered a cell-intrinsic metabolic divergence contributing to the markedly slower pace of human development. GPD1L, a glycerol-3-phosphate dehydrogenase, emerged as differentially expressed between species, linking NADH redox metabolism to developmental rate. By measuring NAD⁺/NADH ratios and glycolytic flux with orthogonal tracing approaches, we found that human cells exhibit a more reduced redox state than mouse cells, consistent with lower NADH oxidation capacity. Importantly, we restricted our redox measurements to the cytosol using the dedicated SoNar sensor, capturing the compartment-specific role of NAD⁺/NADH in regulating tempo.

Temporal acceleration of developmental processes was achieved by enhancing NADH oxidation: CRISPRa-driven upregulation of GPD1/1L, or ectopic expression of potent oxidoreductases, increased cytosolic NAD⁺ regeneration and accelerated multiple embryonic events *in vitro*. Boosting NAD⁺ levels shortened the segmentation clock period and sped up differentiation into definitive endoderm and neural progenitor cells. This suggests that metabolic tuning can synchronize the tempo of distinct developmental trajectories. The demonstration that modulation of GPD1L accelerates the cell cycle, segmentation clock, endoderm differentiation, and neural fate acquisition suggests a coordinated temporal control mechanism downstream of redox balance. Our findings address a long-standing gap in understanding how developmental timing is controlled across species. Prior work had linked metabolism to tempo regulation: for instance, faster-developing species exhibit higher metabolic rates, and perturbing glycolysis or mitochondrial function can decelerate segmentation(*21, 35*). Yet, these approaches did not identify causal endogenous factors and often relied on broad-acting small molecules with poor specificity. Such interventions frequently lack consideration for long-term developmental outcomes and do not test whether gain- or loss-of-function phenotypes are compatible with embryogenesis. In contrast, our identification of GPD1L offers a defined genetic entry point to control pace without compromising viability. This adds a mechanistic layer to recent insights into how protein degradation(*65*) and metabolic state jointly regulate species-specific tempo. By identifying a single metabolic gene that can tune developmental speed across lineages, our work supports the existence of a “tempo control” module rooted in intracellular redox biochemistry.

Recent work supports a signaling function for metabolism in development(*14*). Glycolytic flux can act independently of energy output to regulate segmentation via Wnt signaling(*14, 15, 61*). This modular architecture of metabolic signaling suggests that GPD1L-mediated redox control may act through parallel or convergent pathways, reinforcing its role as a tempo modulator.

Our data align with multiomic profiling that revealed species-specific regulation of glycogen metabolism in early development(*66*) and with emerging evidence for redox divergence in glycerol-3-phosphate metabolism across mammals(*67*). However, this metabolic configuration appears to be specific to the embryonic state. Human fetal and postnatal tissues display measurable G3P shuttle activity(*44, 68*), indicating that reduced cytosolic NADH oxidation is not a fixed property of the species but rather a transient feature of early development. Thus, tempo control likely reflects an embryogenesis-restricted metabolic wiring that is later remodeled during tissue maturation. These findings point to a broader evolutionary strategy where metabolic rewiring contributes to species-specific timing.

The regulatory function of redox metabolism in developmental tempo is likely conserved beyond mammals. In *Drosophila*, null alleles of *Gpdh1*, the sole *GPD1/L* ortholog, give rise to a lowered NAD^+^/NADH ratio and developmental delay with decreased larval body size. Combined loss of both *Gpdh1* and *Ldh* results in severe growth defects and markedly reduced viability(*69*). A recent study also found that NAD^+^ availability mediates the delay in progression of a differentiation front in the developing *Drosophila* eye in the context of electron transport chain deficiency(*24*). The connection between developmental rate and redox metabolism may therefore be deeply conserved.

From a translational perspective, targeted manipulation of NADH metabolism could accelerate organoid maturation, align cross-species developmental benchmarks, or enhance differentiation protocols. Even modest acceleration of human stem cell differentiation could improve disease modeling or therapeutic testing. More broadly, our findings establish metabolism as an evolutionary dial for developmental rate. The ability to reprogram intrinsic metabolic circuits to shift the pace of embryonic processes provides a conceptual and molecular framework for understanding and engineering the tempo of human development.

## Acknowledgments

Feedback on the manuscript was provided by members of the Diaz Cuadros laboratory, Konrad Hochedlinger, Radhika Subramanian, Owen Skinner, Fred Ausubel, Olivier Pourquié, Gary Ruvkun and Michael Duchen. We thank members of the Gary Ruvkun and Jonathan Strecker laboratories for helpful discussions. Human iPS and mouse ES cell lines were provided by the Olivier Pourquié laboratory. Mouse EpiSCs were provided by Jun Wu, the PAX6::GFP line by Lorenz Studer, and the RUES2-GLR line by Ali Brivanlou. The miRFP670 construct was provided by Michael Kossifos. Confocal microscopy was performed with assistance from Sam Wattrus using equipment in his laboratory. Cell sorting was performed at the Center for Computational and Integrative Biology (CCIB) at Massachusetts General Hospital using a SONY SH800 instrument. Proteomic analyses were performed at the Thermo Fisher Scientific Center for Multiplexed Proteomics (TCMP) at Harvard Medical School (https://tcmp.hms.harvard.edu). Seahorse assays were performed using the XFe96 instrument in Vamsi Mootha’s laboratory. VKM is an Investigator of the Howard Hughes Medical Institute.

## Funding

Research in the Diaz Cuadros lab was funded by the Department of Molecular Biology at Massachusetts General Hospital, the Physician and/or Scientist Development Award from the Massachusetts General Hospital Executive Committee on Research, by the Eunice Kennedy Shriver National Institute of Child Health and Human Development (NICHD) of the National Institutes of Health (NIH) under grant number R01HD120311, and partially by the National Institute of General Medical Sciences (NIGMS) of the National Institutes of Health (NIH) under grant number R01GM153752. The content is solely the responsibility of the authors and does not necessarily represent the official views of the National Institutes of Health.

## Author contributions

G.E.V. and M.D.C. conceived the project, designed the experiments, performed the research, and wrote the manuscript with input from all authors. X.Z. developed the bioinformatic pipelines for scRNA-seq analysis. C.M., L.A., D.A., KI and J.P.M. assisted in the design and generation of cell lines used in this study. J.P.M, L.A., S.C. and C.M. also collaborated in the extraction and preparation of omics samples. L.A. collaborated in the recording and analysis of FUCCI data. H.Z., Y.W., and G.P. performed and analyzed the deuterium-tracing experiments. F.D., R.S., and V.M. performed and analyzed the 13C-glucose metabolomics. M.D.C supervised the project.

## Competing interests

G.J.P. is the Chief Scientific Officer of Panome Bio and a scientific advisory board member for Cambridge Isotope Laboratories. The Patti laboratory has a collaborative agreement with Thermo Fisher Scientific. VKM is an advisor to and receives equity from 5am Ventures and Falcon Bio. All other authors declare no competing interests.

## Data, code, and materials availability

the datasets generated during and/or analyzed during the current study are available from the corresponding authors upon request. High throughput sequencing data generated in this study have been deposited in the NCBI Gene Expression Omnibus under accession number **GSE310911**. All materials used in this study, including stem cell lines carrying knock-in reporters, are available by request from the corresponding author.

All custom code used for re analyzing the single cell RNA sequencing datasets is available on GitHub at: https://github.com/gesvaldebenito-gif/redoxregulation. The repository includes the full R scripts for R Studio using Seurat v5 in R Version 2024.09.1+394 and provides all steps required to reproduce the analyses presented in this study.

## Materials and Methods

### Culture and maintenance of cell lines

Mouse epiblast stem cells (EpiSCs) were obtained from the laboratory of Jun Wu (University of Texas Southwestern Medical Center) and cultured under feeder-dependent conditions in NFBR + 10% KSR medium, freshly prepared each day(*70*). NFBR medium consisted of N2B27 base medium composed of DMEM/F12 (Thermo Fisher, cat. no. 11320033), Neurobasal (Thermo Fisher, cat. no. 21103049), 1× N2 Supplement (Thermo Fisher, cat. no. 17502048), 1× B27 Supplement (Thermo Fisher, cat. no. 17504044), 1× GlutaMAX (Thermo Fisher, cat. no. 35050061), 1× non-essential amino acids (NEAA; Thermo Fisher, cat. no. 11140050), and 0.1 mM β-mercaptoethanol (Thermo Fisher, cat. no. 21985-023). The base was supplemented with 10% KnockOut Serum Replacement (Thermo Fisher, cat. no. 10828-028), 20 ng mL⁻¹ FGF2 (PeproTech, cat. no. 450-33), 2.5 µM IWR-1 (Sigma-Aldrich, cat. no. I0161), and 1 mg mL⁻¹ BSA (Sigma-Aldrich, cat. no. A7030). Cultures were maintained on mitotically inactivated mouse embryonic fibroblast feeders (EMD Millipore, cat. no. PMEF-CF).

To adapt EpiSCs to feeder-free conditions, cells were transferred to matrigel-coated plates (Corning, cat. no. 354277) and cultured in mTeSR Plus medium (STEMCELL Technologies, cat. no. 100-0276) supplemented with 2.5 µM IWR-1 (Sigma-Aldrich, I0161) and 12.5 ng mL⁻¹ Activin A (R&D Systems, cat. no. 338-AC-050) for two weeks. Thereafter, Activin A was withdrawn, and cells were expanded and cryopreserved for subsequent use. Upon thawing, cells were maintained in mTeSR Plus medium without supplements and passaged every two days using ReLeSR (STEMCELL Technologies, cat. no. 100-0484).

Human pluripotent stem cells (hPSCs) were maintained on Matrigel-coated tissue-culture dishes (Corning, cat. no. 354277) in mTeSR Plus medium (STEMCELL Technologies, cat. no. 100-0276). All human embryonic stem cell work was approved by the Massachusetts General Hospital Embryonic Stem Cell Research Oversight (ESCRO) Committee (ESCRO #2022E05_05; IBC #2022B000033). All work with human cell lines was approved by the Massachusetts General Hospital Institutional Review Board (Protocol #2022P000297 – Non-Human Subjects Research determination). We complied with all relevant ethical regulations. Cells were passaged every 4–5 days using ReLeSR (STEMCELL Technologies, cat. no. 100-0484) and maintained at 37 °C in a humidified incubator with 5% CO₂. Human reporter lines used in this study included NCRM1 *HES7-Achilles*(*4*) and hiPS11-a *MSGN1-Venus*(*71*), both obtained from the laboratory of Olivier Pourquié (Harvard Medical School). Additional lines included H9 *PAX6::GFP*, generated in the laboratory of Lorenz Studer (Sloan Kettering Institute, Memorial Sloan Kettering Cancer Center)(*63, 72*), and *RUES2-GLR*, obtained from the laboratory of Ali Brivanlou (The Rockefeller University)(*64*). All human lines were cultured on Matrigel-coated dishes in mTeSR Plus medium under the same CO₂ and temperature conditions described above.

Naive E14 mouse embryonic stem cells carrying the *pMsgn1-Venus*(*32*) reporter were obtained from Olivier Pourquié and maintained in 2i media as previously described(*21*).

All cells were routinely tested for mycoplasma contamination using the MycoStrip™ mycoplasma detection kit (InvivoGen, cat. no. rep-mys-50).

A summary of all cell lines used in this study is provided in **Table S1**.

### Generation of mouse EpiSC reporter lines

Mouse EpiSC *Msgn1-Achilles* and *Hes7-Achilles* reporter knock-in lines were generated by CRISPR-Cas9-mediated homology-directed repair. Single-guide RNAs were designed to target the 3′ end of each locus (immediately upstream of the stop codon) using the CRISPOR design tool and cloned into pGuide-it-tdTomato (Takara, cat. no. 632604). For *Hes7*, we implemented the same targeting strategy previously used for naive embryonic stem cells(*4*). Briefly, the donor plasmid consisted of 1-kb 5′ and 3′ homology arms flanking a cassette encoding T2A–Achilles– NLS–CL1–PEST in a pUC19 backbone assembled by Gibson cloning; the PAM sequence in the donor was mutated by site-directed mutagenesis (In-Fusion, Takara) to prevent re-cleavage. For *Msgn1*, the donor plasmid carried 1-kb 5′ and 3′ homology arms flanking T2A–Achilles– NLS (no destabilization domains). The Achilles fragment was subcloned from the pHes7-Achilles-Hes7 plasmid (Addgene cat. no. 153528).

Plasmids (pGuide-it vector and donor vector) were introduced into mouse EpiSCs cultured in mTeSR Plus media on matrigel-coated dishes using Lipofectamine Stem (Invitrogen, cat. no. STEM00003). Twenty-four hours later, tdTomato⁺ cells were enriched by flow cytometry (SONY SH800) and seeded at low density on matrigel-coated dishes for clonal expansion. Individual colonies were expanded, genotyped by PCR for correct insertion at the 3′ end of *Hes7* (T2A– Achilles–NLS–CL1–PEST) or *Msgn1* (T2A–Achilles–NLS), and verified by Sanger sequencing to exclude undesired mutations at the targeted loci. Positive clones were banked and retested after thawing.

A summary of all cell lines used in this study is provided in **Table S1**.

### Directed differentiation

For presomitic mesoderm induction, mouse EpiSCs and human pluripotent stem cells were seeded at densities of 1.5 × 10⁵ cells per cm² (mouse) and 2.5 × 10⁵ cells per cm² (human) on Matrigel-coated plates (Corning, cat. no. 354277) and cultured in mTeSR Plus medium (StemCell Technologies, cat. no. 100-0276). Human cells were supplemented with 10 µM Y-27632 (Tocris, cat. no. 1254) to enhance survival. After 24 hours, the medium was replaced with fresh mTeSR Plus without Y-27632.

The following day (day 0 of differentiation), cultures were switched to DiCLF medium, consisting of DMEM/F12 (Thermo Fisher, cat. no. 11320033) supplemented with 1× ITS (Thermo Fisher, cat. no. 41400045), 6 µM CHIR99021 (Tocris, cat. no. 4423/10), 0.5 µM LDN-193189 (Axon Medchem, cat. no. AMS.04-0074), and 20 ng mL⁻¹ FGF2 (PeproTech, cat. no. 450-33). Medium was refreshed daily throughout the differentiation period.

For experiments requiring extended maintenance of the presomitic mesoderm (PSM) state, cells were cultured in Extension medium, composed of DMEM/F12 supplemented with 1% ITS, 10% KnockOut Serum Replacement (Thermo Fisher, cat. no. 10828-028), 2 µg mL⁻¹ Heparin (Sigma-Aldrich, cat. no. H3393-100KU), 6 µM CHIR99021 (Tocris, 4423/10), 0.5 µM LDN-193189 (Axon Medchem, AMS.04-0074), 20 ng mL⁻¹ FGF4 (R&D Systems, cat. no. 5846-F4-025), 2.5 µM BMS493 (Sigma-Aldrich, cat. no. B6688-5MG), and 10 µM Y-27632 (Tocris, cat. no. 1254/10).

For time-lapse experiments, cells were seeded on Matrigel-coated ibidi imaging plates (ibidi, cat. no. 80636) and cultured in Extension medium in which DMEM/F12 was replaced with FluoroBrite DMEM (Thermo Fisher, cat. no. A1896702) supplemented with 1× GlutaMAX (Thermo Fisher, cat. no. 35050061), 1× non-essential amino acids (NEAA; Thermo Fisher, cat. no. 11140050), and 1 mM pyruvate (Gibco, cat. no. 11360070), together with 1% ITS, 10% KnockOut Serum Replacement (Thermo Fisher, cat. no. 10828-028), 2 µg mL⁻¹ heparin (Sigma-Aldrich, cat. no. H3393-100KU), 6 µM CHIR99021 (Tocris, 4423/10), 0.5 µM LDN-193189 (Axon Medchem, AMS.04-0074), 20 ng mL⁻¹ FGF4 (R&D Systems, cat. no. 5846-F4-025), 2.5 µM BMS493 (Sigma-Aldrich, cat. no. B6688-5MG), and 10 µM Y-27632 (Tocris, cat. no. 1254/10).

Naive mouse embryonic stem cells were pre-differentiated to epiblast-like fate and subsequently differentiated to presomitic mesoderm as previously described(*21*).

For definitive endoderm differentiation, human pluripotent stem cells were seeded at 1.0 × 10⁵ cells per cm², and mouse epiblast stem cells at 0.8 × 10⁵ cells per cm², on Matrigel-coated plates (Corning, cat. no. 354277) in mTeSR Plus medium (STEMCELL Technologies, cat. no. 100-0276) supplemented with 10 µM Y-27632 (Tocris, cat. no. 1254/10) for 24 hours. The medium was then replaced with DMEM (1×) containing 10 ng mL⁻¹ Activin A (R&D Systems, cat. no. 338-AC-050), 3 µM CHIR99021 (Tocris, cat. no. 4423/10), 10 ng mL⁻¹ FGF2(PeproTech, cat. no. 450-33), and 1 mM sodium pyruvate (Thermo Fisher, cat. no. 11360070). Medium was refreshed daily for the duration of the induction.

For anterior neuroectoderm differentiation, human pluripotent stem cells were seeded at 4.0 × 10⁵ cells per cm², and mouse epiblast stem cells at 3.5 × 10⁵ cells per cm², on Matrigel-coated plates (Corning, cat. no. 354277) in mTeSR Plus medium (STEMCELL Technologies, cat. no. 100-0276) supplemented with 10 µM Y-27632 (Tocris, cat. no. 1254/10) for 24 hours. The medium was then replaced with fresh mTeSR Plus lacking Y-27632. The following day, cultures were switched to dual-SMAD inhibition medium, consisting of N2B27 supplemented with 10 µM SB431542 (Selleckchem, cat. no. S1067) and 100 nM LDN-193189 (Axon Medchem, cat. no. AMS.04-0074). Medium was refreshed daily, and cells were maintained at 37 °C, 5% CO₂ in a humidified incubator until analysis.

### Time-lapse imaging

Time-lapse imaging was performed on a Leica DMi8 inverted fluorescence microscope equipped with a temperature- and CO₂-controlled incubation chamber (37 °C, 5% CO₂) using a 20× objective. For Achilles, excitation was at 510 nm with a 535/50 nm emission filter; images were acquired with a 1 s exposure at 75% light intensity. For Venus, excitation was at 475 nm with a 535/50 nm emission filter; images were acquired with a 100 ms exposure at 30% light intensity. Imaging intervals were as follows: HES7–Achilles was imaged every 15 min for 48 h, MSGN1– Venus every 6 h for mouse–human comparative experiments, and every 1 h for overexpression experiments.

### Image processing and oscillation analysis

All fluorescence time-lapse sequences were background-subtracted in Fiji to enhance signal-to-noise ratio. For quantification of Achilles oscillations, small regions of interest (ROIs) were drawn over the reporter-positive areas, and the mean fluorescence intensity over time was extracted. Intensity profiles were smoothened in GraphPad Prism using six neighboring data points and a second-order polynomial for visualization purposes. For period quantification, peaks were identified manually in the raw (non-smoothened) traces, and the time intervals between consecutive peaks were used to calculate oscillatory periods. The segmentation clock period corresponds to the mean time between two HES7–Achilles peaks across individual traces from the same experimental batch. Representative traces shown in figures correspond to mean ± s.e.m. of ROI-based profiles, normalized in GraphPad Prism such that the smallest mean intensity in each dataset was set to 0% and the largest mean to 100%.

### Single-cell oscillation tracking

For single-cell analyses, HES7–Achilles reporter cells were co-infected with a constitutive nuclear label (H2B–mCherry; Addgene, cat. no. 21217) to enable reliable nuclear segmentation and longitudinal tracking. Cell tracking was performed in Fiji using the TrackMate plugin with the StarDist detector to identify individual nuclei. Simple LAP tracker parameters were linking max distance: 40 pixels, gap-closing max distance: 20 pixels, and gap-closing max frame gap: 2. Tracks were inspected and manually corrected to ensure continuous single-cell trajectories across all frames. For each tracked cell, the mean Achilles fluorescence intensity within the nuclear ROI was plotted as a function of time to generate individual oscillation curves.

### MSGN1–Venus reporter induction analysis

MSGN1–Venus reporter cells also carried a constitutive nuclear label (H2B–mCherry; Addgene, cat. no. 21217), which was used for accurate nuclear segmentation and motion vector (MV) quantification. Nuclear centroids were detected in each frame using the StarDist plugin. A cell was considered MSGN1-positive once its Venus fluorescence intensity exceeded a predefined threshold above background, applied consistently across all time-lapse fields. MV % was defined as the proportion of MSGN1-positive cells relative to the total number of H2B–mCherry–labeled cells.

### PAX6::GFP and SOX17::tdTomato reporter induction analyses

Cells were plated in ibidi plates (ibidi, cat. no. 80636) and differentiated as described above. Imaging was performed on a Nikon CSU-W1 spinning-disk confocal microscope equipped with a Prime BSI sCMOS camera, a 40×/1.15 NA objective, and an environmental chamber maintained at 37 °C and 5 % CO₂. For PAX6::GFP, excitation was achieved with the 488 nm laser (50 % power) and emission was collected using a GFP filter (approximately 525/50 nm). For SOX17::tdTomato, excitation was performed with the 561 nm laser (20 % power) and emission was collected using a tdTomato filter (approximately 600/50 nm). Images were acquired in 16-bit mode using 2×2 binning and HDR gain, with exposure times set to 6 s for PAX6::GFP and 500 ms for SOX17-tdTomato. Fluorescence intensity was quantified in Fiji after background subtraction.

### Lentivirus production and transduction

Lentiviral particles were generated in HEK293T cells by co-transfecting the packaging plasmids psPAX2 (Addgene cat. no. 12260) and pVSV-G (Addgene cat. no. 12259) together with the desired transfer vector using Lipofectamine 3000 (Thermo Fisher Scientific, cat. no. L3000001), following the manufacturer’s instructions. Transfection mixtures were incubated for 30 min at room temperature before being added to cells cultured in DMEM supplemented with 10% FBS (GeminiBio, cat. no. 900-208). After overnight incubation, the medium was replaced, and viral supernatants were collected 48 h post-transfection. Viral particles were concentrated using Lenti-X Concentrator (Takara, cat. no. 631232) according to the manufacturer’s protocol. For transduction, concentrated viral particles were added to stem cells maintained in mTeSR™ Plus medium (STEMCELL Technologies, cat. no. 100-0276) supplemented with penicillin– streptomycin (Thermo Fisher Scientific, cat. no. 15140122). Depending on the transfer vector, successfully transduced cells were either selected using the appropriate antibiotic resistance marker or purified by fluorescence-activated cell sorting prior to downstream analyses.

### Cloning, validation, and imaging of SoNar construct

The cytosolic NAD⁺/NADH biosensor SoNar (FR Biotechnology) was subcloned from the pcDNA backbone into the lentiviral expression vector pLX_311_KRAB-dCas9 (Addgene, cat. no. 96918). The backbone was linearized by BamHI and NheI digestion to remove the KRAB-dCas9 fragment, and the SoNar coding sequence was inserted using Gibson assembly (New England Biolabs cat. no. E5520S) following the manufacturer’s protocol. The resulting construct, pLX_311-SoNar, places the sensor under the EF1α promoter and confers blasticidin resistance for stable selection.

Lentivirus was produced as described above and used to transduce pluripotent stem cells, which were subsequently selected with blasticidin until a pure population of SoNar-positive cells was established.

Live-cell imaging was performed on a Nikon CSU-W1 spinning-disk confocal microscope equipped with a Prime BSI sCMOS camera, a 40×/1.15 objective, and an environmental chamber maintained at 37 °C, 5 % CO₂, and 5 % O₂. The SoNar ratiometric signal was recorded using 405 nm and 488nm laser excitation lines with corresponding emission collection at 450–490 nm and 510–550 nm, respectively. Images were acquired sequentially using NIS-Elements ND Acquisition with exposure times of 150 ms per channel. The fluorescence ratio (488/405nm) was calculated in FIJI after background subtraction to represent the relative NAD⁺/NADH redox state.

To validate biosensor performance, cells were first incubated in glucose-free medium (Thermo Fisher Scientific, cat. no. A1443001), followed by sequential addition of 17.5 mM D-glucose and 1 mM sodium pyruvate (Gibco, cat. no. 11360070), which produced the expected reciprocal changes in the excitation ratio, confirming sensor responsiveness to dynamic cytosolic NAD⁺/NADH fluctuations.

### *Lb*NOX and *Ec*STH constructs

The coding sequences for *Lb*NOX (*Lactobacillus brevis NADH oxidase*)(*40*) and *Ec*STH (*Escherichia coli soluble transhydrogenase*)(*42*) were subcloned into the lentiviral expression vector pLX_311_KRAB-dCas9 (Addgene, cat. no. 96918) as described above for SoNar. The *Lb*NOX fragment was obtained from *pUC57-LbNOX* (Addgene cat. no. 75285) and *Ec*STH from *pEcSTH* (Addgene cat. no. 211923). Each construct was designed to co-express miRFP670 (Addgene cat. no. 79987) polycistronically using a T2A peptide cloned after the *Lb*NOX or *Ec*STH coding sequences to enable visualization of transduced cells. An empty vector control expressing miRFP670 only was generated in parallel. Lentivirus was produced and used for stem cell transduction as described above. Following infection, cells were selected with blasticidin until a stable population was obtained. Successful expression was confirmed by miRFP670 fluorescence using an EVOS imaging system (Thermo Fisher Scientific) equipped with the Cy5 filter cube.

### Analysis of published single-cell RNA sequencing data

Single-cell RNA-seq datasets from the human PCW3 embryo(*51*), E8.0 mouse embryo(*50*), and *in vitro* differentiation system(*4*) were analyzed using Seurat v5 in R Studio. Raw 10x Genomics count matrices were imported with Read10X and converted into Seurat objects. Low-quality cells were excluded based on standard thresholds (n_features < 200 or > 6,000, n_counts > 50,000, or >10% mitochondrial transcripts). Counts were log-normalized, and the top 2,000 highly variable genes were selected. Data were scaled and regressed for mitochondrial content and UMI counts, followed by principal component analysis (PCA), nearest-neighbor graph construction, Louvain clustering (resolution = 0.2), and UMAP embedding using the first 20–30 principal components. Cluster marker genes were identified using the Wilcoxon rank-sum test (FindAllMarkers, min.pct = 0.1, logFC > 0.15) and exported for downstream annotation. Clusters were assigned to ectoderm, mesoderm, or endoderm identities based on canonical lineage markers. Gene-level expression was visualized using FeaturePlot and DimPlot functions.

### Bulk RNA sequencing and analysis

Cells were differentiated as described above, and total RNA was extracted using TRIzol Reagent (Thermo Fisher Scientific, cat. no. 15596026) following the manufacturer’s protocol. RNA concentration and purity were determined by Qubit and NanoDrop, and integrity was confirmed using an Agilent 2100 Bioanalyzer (RIN ≥ 4.0). Library preparation and sequencing were performed by Novogene (Sacramento, USA). For both human and mouse samples, poly(A)-enriched mRNA libraries were generated and sequenced on an Illumina NovaSeq X Plus platform using paired-end 150 bp reads, generating approximately 6 Gb of raw data per sample.

Data processing included adapter trimming and quality filtering, read alignment to the respective reference genomes, and quantification of gene-level expression counts. For intraspecies analyses, raw counts were used directly. For interspecies comparisons, counts were normalized as fragments per kilobase per million (FPKM), and only one-to-one orthologous genes between human and mouse were retained. Orthology was determined using Ensembl BioMart (release 52) by selecting “Homo sapiens genes” and filtering for “orthologous mouse genes.” Non-homologous genes were excluded prior to downstream differential expression and pathway enrichment analyses.

Data visualization, including heat maps, volcano plots, and dot plots, was performed in GraphPad Prism version 10.5.0 (build 673).

### Differential expression and gene set analysis

Differential expression and clustering analyses were performed using iDEP 2.01 (https://bioinformatics.sdstate.edu/idep/). Normalized expression data (FPKM) were used for interspecies comparisons, while raw counts were used for intraspecies analyses. Genes with FPKM values greater than 1 in at least three samples were retained. Hierarchical clustering was performed using Pearson correlation and average linkage, typically selecting the top 500–1000 most variable genes. For differential expression, the limma-trend method was applied to normalized expression data, and the limma-voom approach was used for raw count data to account for mean–variance relationships. Significance was defined at adjusted p < 0.05. Heatmaps were generated using gene-wise z-score scaling to emphasize relative expression differences across samples. Gene Ontology and KEGG pathway enrichment analyses were performed on significant gene sets to identify overrepresented biological processes.

### Proteomic sample preparation and analysis

Cells were differentiated as described above, and proteins were extracted using RIPA buffer (Sigma-Aldrich, cat. no. R0278) by incubating samples for 30 minutes on ice. Following extraction, protein lysates were snap-frozen in liquid nitrogen and submitted to the Thermo Fisher Scientific Center for Multiplexed Proteomics at Harvard Medical School for mass spectrometry-based quantitative analysis. Multiplexed proteomic profiling was performed using high-resolution TMT mass spectrometry. Final data were normalized to the sum of signal-to-noise ratios for all detected proteins and expressed as scaled relative abundance values.

### ^13^C-glucose tracing, metabolite extraction, and LC-MS

Cells were seeded and differentiated into presomitic mesoderm (PSM) as described above. On day 2 of differentiation, cultures were incubated for 24 h in Extension medium prepared using glucose-free DMEM (Gibco cat. No. 11966025) supplemented with 1× GlutaMAX (Thermo Fisher, cat. no. 35050061), 1× non-essential amino acids (NEAA; Thermo Fisher, cat. no. 11140050), 1 mM pyruvate (Gibco, cat. no. 11360070) and either labeled U-^13^C_6_-D-glucose (Cambridge Isotope Laboratories, cat. no. CLM-1396) or unlabeled D-glucose at a final concentration of 17.5 mM. After incubation, cells were washed twice with ice-cold phosphate-buffered saline (PBS) and immediately frozen in liquid nitrogen. Metabolites were extracted on ice by adding pre-chilled extraction solvent of 4:4:2 MeOH/ACN/H_2_O (v/v/v) containing 0.1 M formic acid to each well, followed by cell scraping and transferring to microcentrifuge tubes. Samples were vortexed for 10 s, incubated on ice for 3 min, neutralized with 15% ammonium bicarbonate, and placed on dry ice for 20 min. Tubes were centrifuged at 21,000 × g for 20 min at 4 °C, and supernatant was transferred to LC-MS vials for further analysis.

The separation was performed using an Xbridge BEH Amide column (2 mm × 150 mm × 2.5 µm; Waters, Milford, MA) at a column temperature of 25°C. Solvent A consisted of 5% acetonitrile (v/v) and 20 mM ammonium acetate at pH 9.0, while solvent B was 100% acetonitrile. The gradient program was as follows: 0 min, 90% B; 2 min, 90% B; 3 min, 75% B; 7 min, 75% B; 8 min, 70% B; 9 min, 70% B; 10 min, 50% B; 12 min, 50% B; 13 min, 25% B; 14 min, 25% B; 16 min, 0% B; 20.5 min, 0% B; 21 min, 90% B; 25 min, 90% B. The total run time was 25 min with a flow rate of 0.15 mL/min. The Q Exactive Plus mass spectrometer (Thermo Fisher Scientific, Waltham, MA) operated in polarity switching mode, with a scan range of m/z 70–1000 (negative mode) and m/z 120–1000 (positive mode), and a resolution of 140,000 at m/z 200. Additional MS parameters were as follows: sheath gas flow rate 50, auxiliary gas flow rate 10, sweep gas flow rate 2, spray voltage 3.3 kV, capillary temperature 310°C, S-lens RF level 50, AGC target 3E6, and maximum injection time 200 ms.

Nucleotide analysis was performed using a ZIC-pHILIC column (2.1 mm × 150 mm × 5 µm). Buffer A was 20 mM ammonium carbonate, pH 9.6 and buffer B was 100% acetonitrile. The total flow rate was set to 0.15 mL/min and the samples were loaded at 80% B. The gradient was maintained at 80% B for 0.5 min, then ramped to 20% B over the next 20 min, held at 20% B for 0.8 min, ramped to 80% B over 0.2 min, then held at 80% B for 7.5 min for re-equilibration. The elute was directed into Q-Exactive Plus Orbitrap mass spectrometer (Thermo Fisher Scientific, Waltham, MA) with HESI probe operating in polarity switching mode. MS parameters were: sheath gas flow 30, aux gas flow 7, sweep gas flow 2, spray voltage 3.3 kV in both negative and positive modes, capillary temperature 310°C, S-lens RF level 50 and aux gas heater temperature 370°C. Other MS parameters included: resolution of 140,000 at m/z 200, automatic gain control (AGC) target at 1E6, maximum injection time of 80 ms and scan range of m/z 70–1000.

NAD(P)(H) was detected using a 15-minute method with the same LC buffers and column, following this gradient: 0 min, 80% B at a flow rate of 0.25 mL/min; 0.5 min, 80% B at 0.25 mL/min; 8.5 min, 30% B at 0.25 mL/min; 10.5 min, 30% B at 0.25 mL/min; 10.8 min, 80% B at 0.25 mL/min; 11.9 min, 80% B at 0.25 mL/min; 12 min, 80% B at 0.3 mL/min; 14.5 min, 80% B at 0.3 mL/min; 14.7 min, 80% B at 0.25 mL/min; and 15 min, 80% B at 0.25 mL/min. A SIM scan was performed in negative mode, with m/z 620-680 for NAD(H) and m/z 720-780 for NADP(H), with a resolution of 140,000 at m/z 200.

Data were acquired using Xcalibur (v.4.1.31.9, Thermo Fisher). Data were analyzed via MAVEN software using an in-house library(*73, 74*) with 5 ppm mass tolerance and isotope labeling data were corrected for natural isotope abundance (*75*).

### 4-²H-glucose tracing, metabolite extraction, and LC-MS

Cells were seeded and differentiated into PSM as described above. On day 2 of differentiation, cultures were incubated in DiCL medium containing [4-²H]-D-glucose (Cambridge Isotope Laboratories, cat. no. DLM-1278) or unlabeled D-glucose at a final concentration of 17.5 mM as described above. After incubation, cells were collected at 0, 1, 3, 5, 10, 60, and 90 minutes following medium exchange. At each time point, the medium was aspirated, cells were rapidly washed once with ice-cold phosphate-buffered saline (PBS), and metabolism was quenched by addition of ice-cold methanol.

Cells were scraped into methanol, pelleted by centrifugation, and dried by using a SpeedVac concentrator. Dried pellets were extracted with 1 mL methanol/acetonitrile/water (2:2:1, v/v/v), vortexed for 30 s, flash-frozen in liquid nitrogen for 1 min, and sonicated for 10 min. Samples were then incubated at −20 °C for 1 h, centrifuged at 21,000 × g for 20 min at 4 °C, and the supernatant was collected. Extracts were dried by using a SpeedVac and stored at −80 °C until LC/MS analysis.

Cell extracts were resuspended in 250 µL methanol/acetonitrile/water (2:2:1, v/v/v) and analyzed by using a Thermo Scientific Vanquish Horizon UHPLC system coupled to a Thermo Scientific Q Exactive Plus Orbitrap mass spectrometer (Thermo Fisher). The injection volume was 10 µL. Solvent A consisted of 20 mM ammonium bicarbonate, 2.5 µM medronic acid, and 0.1% ammonium hydroxide in water:acetonitrile (95:5), and solvent B consisted of water:acetonitrile (5:95). Metabolites were separated on an iHILIC-(P)-Classic column (100 × 2.1 mm, 5 µm; HILICON). The column was maintained at 40 °C with a flow rate of 0.25 mL min⁻¹. The linear gradient for solvent B was as follows: 0 min, 90%; 1 min, 90%; 14 min, 25%; 14.5 min, 25%; 15 min, 90%; 16.5 min, 90%. The column was equilibrated at 90% B for 4 min at 0.4 mL min⁻¹, followed by 1.5 min at 0.25 mL min⁻¹.

The mass spectrometer, equipped with a heated electrospray ionization source, was operated in negative mode. Key parameters were as follows: ionization voltage, –2.5 kV; sheath gas pressure, 45 arbitrary units (Arb); auxiliary gas, 10 Arb; sweep gas, 2 Arb; auxiliary gas heater temperature, 350 °C; and capillary temperature, 250 °C. Data were acquired in full-scan mode with an *m/z* range of 67–900 at 140,000 resolution, an AGC target of 1 × 10⁶, and a maximum injection time of 200 ms. Data processing and initial annotations were performed by using Xcalibur 4.1 (Thermo Fisher). The identity of a subset of metabolites was confirmed by matching accurate mass, retention time, and MS/MS data to authenticated standards.

The strategy for using [4-²H] glucose to infer shuttle fluxes was described in detail previously (*43*). In brief, relative MDH1 activity was calculated as the fraction of [2-²H] malate (M+1 malate) normalized to the fraction of [1-²H] GAP (M+1 GAP). Relative GPD1/GPD1L activity was calculated as the fraction of [1,2-²H] G3P normalized to the fractions of [1-²H] DHAP and [4-²H] NADH.

GAP and DHAP are structural isomers that cannot be resolved by the LC/MS method used here. We therefore assumed identical labeling fractions for GAP and DHAP, given their rapid interconversion mediated by triosephosphate isomerase. The resolving power of our method also does not distinguish NADH m+1 derived from ^13^C versus ^2^H. To correct for the naturally occurring ^13^C contribution to the deuterated NADH signal, we applied the previously reported computational approach at the following link: https://github.com/e-stan/NADH_KFP.

### Seahorse metabolic flux analysis

Oxygen consumption rate and extracellular acidification rate were measured using the Seahorse XFe96 Analyzer (Agilent). For mitochondrial respiration, the Seahorse XF Cell Mito Stress Test (Agilent, cat. no. 103793-100) was used. Cells were seeded in XF96 cell culture microplates (Agilent, cat. no. 102416-100) and differentiated as described above. On the day of the assay, the culture medium was replaced with Seahorse XF Base Medium (Agilent, cat. no. 103334-100) supplemented with 1 mM pyruvate (Gibco, cat. no. 11360070), 2 mM glutamine (Gibco, cat. no. 25030081), and 10 mM glucose (Gibco, cat. no. A2494001). Plates were incubated for 30 min at 37 °C in a CO₂-free incubator before measurement. Basal respiration was recorded prior to the sequential injection of oligomycin (1 µM; Millipore Sigma, cat. no. 75351-5MG), FCCP (1–2 µM; Millipore Sigma, cat. no. SML2959-1ML), and rotenone/antimycin A (0.5 µM each; rotenone, Millipore Sigma, cat. no. R8875-1G; antimycin A, Millipore Sigma, cat. no. A8674-25MG).

Glycolytic activity was assessed using the Seahorse XF Glycolytic Stress Test Kit (Agilent, cat. no. 103020-100) under similar conditions. Cells were incubated in Seahorse XF Base Medium supplemented with 2 mM glutamine, and ECAR was measured following sequential injection of glucose (10 mM), oligomycin (1 µM; Millipore Sigma, cat. no. 75351-5MG), and 2-deoxy-D-glucose (50 mM; Millipore Sigma, cat. no. D8375). The resulting profiles were used to calculate basal glycolysis, glycolytic capacity, and glycolytic reserve.

To assess complex I–independent respiration, cells were treated with piericidin A and antimycin A by sequential injection, as previously described (*46*) allowing quantification of mitochondrial oxygen consumption driven by alternative electron transport pathways.

After completion of each assay, total protein was extracted from each well, quantified using a BCA assay, and used to normalize OCR and ECAR values.

### Generation of dCas9-VP64 cell lines

Engineered human pluripotent stem cells constitutively expressing transcriptional activator variant of dCas9 were generated by targeted integration into safe-harbor loci. For activation, a *CAG-dCas9-VP64* cassette was integrated into the *AAVS1* locus(*76*). The *dCas9-VP64* fragment was subcloned from the lenti-dCas9-VP64-Blast plasmid (Addgene cat. no. 61425) and inserted in the pAAVS1-Nst-CAG-Dest targeting vector (Addgene cat. No. 80489). Plasmids encoding both Cas9 and an sgRNA targeting either *AAVS1* (pXAT2, Addgene no. 80494) were used. These plasmids and the corresponding donor repair templates were co-transfected using Lipofectamine Stem Transfection Reagent (Thermo Fisher Scientific). After transfection, cells were cultured in mTeSR™ Plus medium (STEMCELL Technologies, cat. no. 100-0276) supplemented with CloneR™ 2 (STEMCELL Technologies, cat. no. 100-0691) to support clonal survival and expansion. Cells were then selected with Geneticin and clonally expanded. Individual clones were validated by PCR to contain the desired insert as previously described(*76*). This process was performed independently in the NCRM1 line, NCRM1 HES7-Achilles line, hiPS11-a MSGN1-Venus line, and H9 PAX6::GFP line.

As a neomycin resistance cassette was already integrated in the RUES2-GLR line, instead of generating a knock-in dCas9–VP64 line, the construct was delivered via lentiviral transduction to achieve random genomic integration using the lenti-dCas9-VP64-Blast transfer vector (Addgene cat. no. 61425). Similarly, the NCRM1 FUCCI line already carried an insert in the AAVS1 locus, so we instead transduced the cells with the lenti-dCas9-VP64-Blast transfer vector (Addgene cat. no. 61425).

### CRISPRa sgRNA design and cloning

Single guide RNAs (sgRNAs) for CRISPR activation (CRISPRa) experiments were designed following established guidelines (*58, 77*). Candidate sgRNAs (**Table S2**) were selected using the CRISPick online tool (Broad Institute; https://portals.broadinstitute.org/gppx/crispick/public). The appropriate reference genome (human or mouse) was selected, and the SpyoCas9 enzyme with the “Chen (2013) tracrRNA” configuration was used as the targeting mechanism. For each target gene, the top-ranked sgRNA was chosen based on on-target efficiency and minimal predicted off-target activity.

sgRNA protospacer sequences were ordered as single-stranded DNA oligonucleotides and cloned into pXPR_502 (CRISPRa; Addgene cat. no. 96923) backbone(*60*). Each sgRNA was first amplified by PCR using universal primers (Amplify_sgRNA_fwd / Amplify_sgRNA_rev, **Table S3**) and Q5 High-Fidelity DNA Polymerase (New England Biolabs), then purified with the DNA Clean & Concentrator Kit (Zymo Research). Purified amplicons were inserted into the target plasmid by Golden Gate assembly using BsmBI-v2 and T4 DNA Ligase (New England Biolabs cat. no. E1602L). Following assembly, plasmids were transformed into chemically competent *E. coli*, individual colonies were expanded in LB with the appropriate antibiotic, and plasmid DNA was isolated by miniprep (QIAGEN). Correct sgRNA insertion was confirmed by Sanger sequencing using a U6 promoter primer.

The resulting transfer vectors were used to produce lentiviral particles and *CAG-dCas9-VP64* cell lines were transduced as described above. Successfully transduced cells were selected with puromycin until a pure population was obtained. Overexpression was validated by immunoblotting or immunofluorescence (see below).

### Generation of FUCCI cell line

The NCRM1 human FUCCI cell cycle reporter line was generated by targeting the AAVS1 safe harbor locus as previously described(*76*) using the pXAT2 (Addgene cat. no. 80494) sgRNA vector and the AAVS1-Puro CAG-FUCCI repair donor vector (Addgene cat. no. 136934)(*78*). These plasmids were co-transfected using Lipofectamine Stem Transfection Reagent (Thermo Fisher Scientific). After transfection, cells were cultured in mTeSR™ Plus medium (STEMCELL Technologies, cat. no. 100-0276) supplemented with CloneR™ 2 (STEMCELL Technologies, cat. no. 100-0691) to support clonal survival and expansion. Cells were then selected with puromycin and clonally expanded. Individual clones were subjected to PCR genotyping as previously described(*76*).

### FUCCI imaging and analysis

Imaging of FUCCI cells was performed on a Leica DMi8 inverted fluorescence microscope equipped with a temperature- and CO₂-controlled incubation chamber (37 °C, 5% CO₂). Cells were seeded on Matrigel-coated ibidi imaging plates (ibidi, cat. no. 80636) and maintained in Extension medium. Time-lapse images were acquired every 30 min for 48 h using the following excitation and emission settings: mCherry, Ex = 635 nm / Em = 642 nm (G1 phase); YFP, Ex = 555 nm / Em = 590 nm (G1/S transition); and GFP, Ex = 510 nm **/** Em = 535 nm (S/G2 phase). Mitosis was identified by characteristic morphological changes such as cell rounding, chromatin condensation, and segregation, visualized using the H2B–iRFP nuclear marker (Ex = 747 nm), which was captured without an emission filter at 100% intensity. The H2B–iRFP construct was generated by lentiviral transduction using the pLenti-H2B-iRFP720 vector (Addgene 128961), which encodes histone H2B fused to iRFP, enabling chromatin labeling in live cells. A complete cell cycle was defined as the time interval from the onset of one mitotic event to the frame immediately preceding the next mitosis.

### Immunoblotting

Cells were washed once with ice-cold phosphate-buffered saline (PBS) and lysed in RIPA buffer (Sigma-Aldrich, cat. no. R0278) supplemented with Protease and Phosphatase Inhibitor Cocktail (Thermo Fisher Scientific, cat. no. 78440). Lysates were scraped and stored at −80 °C until use. Protein concentration was determined using the Pierce BCA Protein Assay Kit (Thermo Fisher Scientific, cat. no. 23227). For immunoblotting, 30 µg of protein per sample was mixed with NuPAGE 4× LDS Sample Buffer (Invitrogen, cat. no. NP0007) and 2% β-mercaptoethanol (Sigma-Aldrich, cat. no. 63689), then boiled at 99 °C for 5 min. Proteins were separated on 4–12% NuPAGE Bis-Tris gels (Invitrogen, cat. no. NP0335) and transferred to Trans-Blot Turbo Mini 0.2 µm Nitrocellulose membranes (Bio-Rad, cat. no. 1704158).

Membranes were blocked for 1 h at room temperature in 2.5% BSA (Sigma-Aldrich, A9418) prepared in DPBS (Gibco, 14190-136), then incubated overnight at 4 °C with primary antibodies (**Table S4**) diluted in blocking buffer containing 0.1% Tween-20 (Sigma-Aldrich, P7949). After washing, membranes were incubated for 1 h at room temperature with IRDye-conjugated secondary antibodies (Li-COR Biosciences; IRDye 680RD Goat anti-Mouse IgG, cat. no. 926-68070; IRDye 800CW Goat anti-Rabbit IgG, cat. no. 926-32211; 1:10,000 dilution). Fluorescent signals were detected using the Odyssey CLx Imaging System (Li-COR Biosciences).

### Immunofluorescence staining

Cells were fixed in 4% paraformaldehyde for 20 min at room temperature, washed three times with phosphate-buffered saline (PBS), and stored in PBS at 4 °C until staining. Fixed cells were permeabilized with 0.1% Tween-20 in Tris-buffered saline (TBS) and blocked for 1 h at room temperature in 3% fetal bovine serum (FBS) and 0.1% Triton X-100 in TBS. Primary antibodies were diluted in blocking solution (1:100–1:1000) and incubated overnight at 4 °C. The following day, cells were washed three times with permeabilization buffer and incubated for 2 h at room temperature in blocking solution containing the corresponding Alexa Fluor–conjugated secondary antibodies (1:500; Thermo Fisher Scientific) and DAPI (1:1000). Samples were washed three times in permeabilization buffer and maintained in PBS prior to imaging.

### Glycerol-3-phosphate (G3P) quantification

Intracellular G3P levels were quantified using the Glycerol-3-Phosphate (G3P) Assay Kit (Colorimetric) (Abcam, cat. no. ab174094) according to the manufacturer’s protocol. Cells were cultured and differentiated as described above, washed twice with ice-cold PBS, and lysed in the assay buffer provided in the kit. Lysates were clarified by centrifugation at 13,000 × g for 10 min at 4 °C, and the supernatant was collected for analysis.

Samples were loaded in triplicate into a 96-well plate along with G3P standards. Reaction mix was added to each well, and plates were incubated for 30–60 min at 37 °C protected from light. Absorbance was measured at 450 nm using a microplate reader. G3P concentrations were determined from the standard curve and normalized to total protein content per well.

### Extracellular lactate quantification

Extracellular lactate levels were measured using the L-Lactate Assay Kit (Colorimetric/Fluorometric, Abcam, cat. no. ab65330) according to the manufacturer’s instructions. Cells were seeded and differentiated into PSM cells as described above. On day 2 of differentiation, cultures were incubated in DiCL medium under standard conditions. At defined time points (0, 1, 3, 5, 10, 60, and 90 minutes), aliquots of conditioned medium were collected and immediately placed on ice. Samples were diluted 1:100 in Assay Buffer to ensure readings within the linear detection range. The colorimetric reaction was developed at room temperature, and absorbance was measured at 570 nm using a microplate reader. Lactate concentrations were calculated from a standard curve and normalized to the basal (time 0) values to account for initial background levels.

### Statistical analyses

Statistical analyses were performed using GraphPad Prism version 10.5.0 (build 673). P values < 0.05 were considered statistically significant. Details of all statistical tests and sample sizes are provided in the corresponding figure legends. Data are presented as mean ± s.d. unless otherwise stated. Unpaired two-tailed Student’s t-tests or ordinary one-way/two-way ANOVA were applied as appropriate, with Tukey’s post-hoc correction for multiple comparisons. All differentiation experiments were performed in at least three independent biological replicates (separate rounds of differentiation), each including three or more technical replicates per condition.

