## Supplementary Data for "A genetically encoded redox bottleneck constrains human developmental rate"

#### Fig. S1.

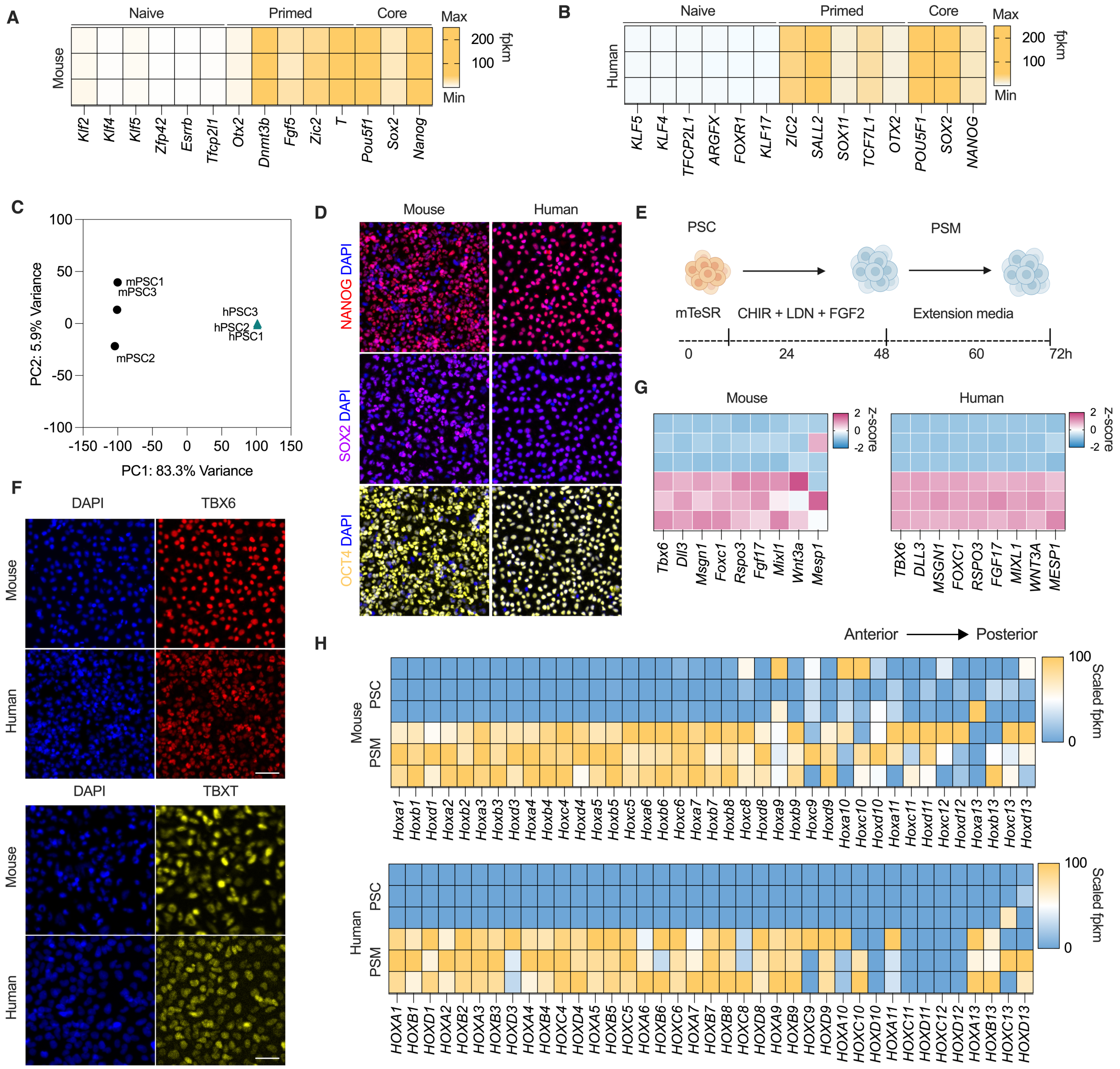

**Figure S1. Validation of pluripotent state and presomitic mesoderm (PSM) differentiation across species.** (**A–B**) Heat maps based on bulk RNA-sequencing data showing normalized expression of pluripotency-associated genes in mouse (**A**) and human (**B**) pluripotent stem cells cultured under identical conditions. Genes are grouped by naïve, primed, and core pluripotency states. (**C**) Principal component analysis (PCA) of bulk RNA-seq data from pluripotent stem cell samples showing clear species segregation along PC1 (83.3% variance). Mouse samples (mPSC; circles) cluster distinctly from human samples (hPSC; triangles), reflecting strong species-specific transcriptional differences. (**D**) Representative immunofluorescence images of pluripotency markers NANOG, SOX2, and POU5F1 (OCT4) in mouse and human pluripotent stem cells prior to differentiation, indicating conserved expression patterns. Scale bar, 100 μm. (**E**) Schematic of the *in vitro* PSM differentiation protocol used for both species, involving mTeSR culture followed by CHIR99021, LDN193189, and FGF2 treatment, then transition to extension medium for 72 h. (**F**) Representative images of TBXT (Brachyury) and TBX6 immunofluorescence at 48 h of differentiation in mouse and human PSM cells, confirming robust mesodermal induction in both mouse and human cultures. Scale bar, 100 μm. (**G**) Heatmap showing expression z-scores for presomitic mesoderm (PSM) marker genes based on bulk RNA-seq profiling of mouse and human undifferentiated pluripotent stem cells and induced PSM cells (n = 3 independent replicates). (**H**) Heat maps based on bulk RNA-sequencing data showing scaled expression of *Hox* genes (mouse *Hoxa1–Hoxd13*, human *HOXA1–HOXD13*) in pluripotent stem cells (PSC) and derived PSM cells, illustrating anterior–posterior patterning along the differentiation axis.

**Fig. S2.**

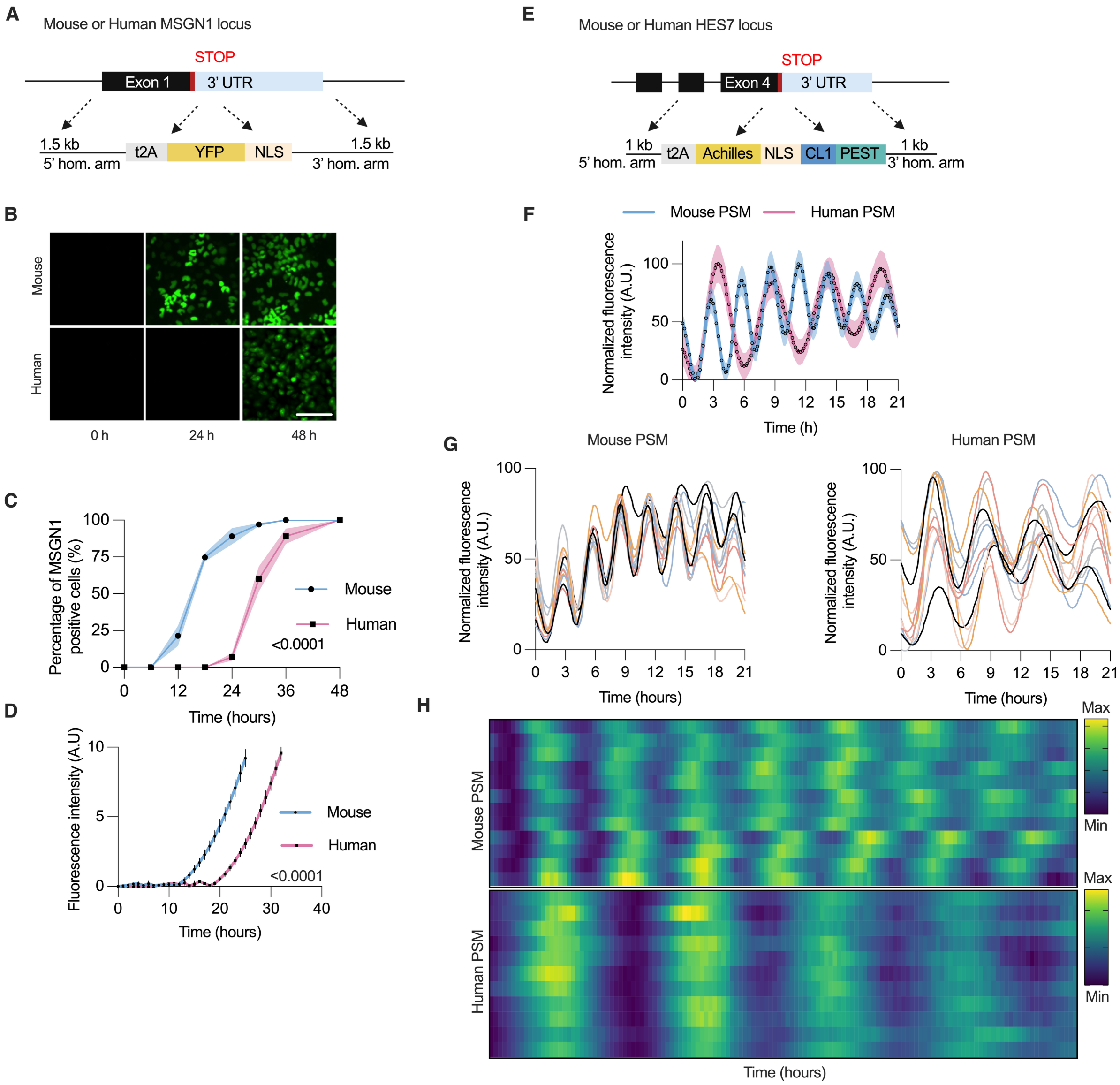

**Fig. S2. Generation and characterization of mouse and human segmentation clock reporter lines.** (**A**) Schematic of the *MSGN1–YFP* reporter design showing targeted insertion of a *T2A–YFP–NLS* cassette immediately upstream of the endogenous stop codon in the mouse *Msgn1* locus and the human *MSGN1* locus. The Achilles YFP protein was used for mouse, whereas the Venus YFP protein was used for human(*71*). (**B**) Representative fluorescence images showing progressive induction of Msgn1-Achilles in differentiating mouse EpiSCs and MSGN1-Venus in differentiating human iPSCs. Day0–Day2 reflect the time elapsed since cultures were placed in differentiation media. Scale bar, 50 µm. (**C**) Quantification of MSGN1-positive cells over time demonstrates faster differentiation kinetics for mouse relative to human PSM, reflecting species-specific tempo differences (n = 3 independent replicates; repeated-measures two-way ANOVA with Tukey correction, $P < 0.0001$). (**D**) Quantification of *MSGN1–YFP* fluorescence intensity during *in vitro* PSM differentiation, showing faster induction kinetics in mouse compared with human cells (n = 3 independent replicates; repeated-measures two-way ANOVA with Tukey correction, $P < 0.0001$). (**E**) Schematic of the *HES7–Achilles* reporter construct(*4*), in which a *T2A–Achilles–NLS–CL1–PEST* cassette was inserted in-frame before the *HES7* stop codon, enabling real-time monitoring of oscillatory dynamics. (**F**) Oscillatory Hes7–Achilles reporter activity recorded in pluripotent stem cell-derived mouse and human PSM cells (n = 3 independent replicates; mean ± s.e.m.). (**G**) Representative *HES7–Achilles* oscillation traces from individual cells in mouse (left) and human (right) pluripotent stem cell-derived PSM cultures, showing shorter oscillation periods in mouse (n = 3 independent experiments per species). (**H**) Heat maps of normalized Achilles fluorescence intensity over time illustrating synchronized *HES7* oscillations in mouse (top) and human (bottom) reporter lines differentiated *in vitro* to PSM fate, with a two-fold difference in period length. Each row represents an individual cell.

**Fig. S3.**

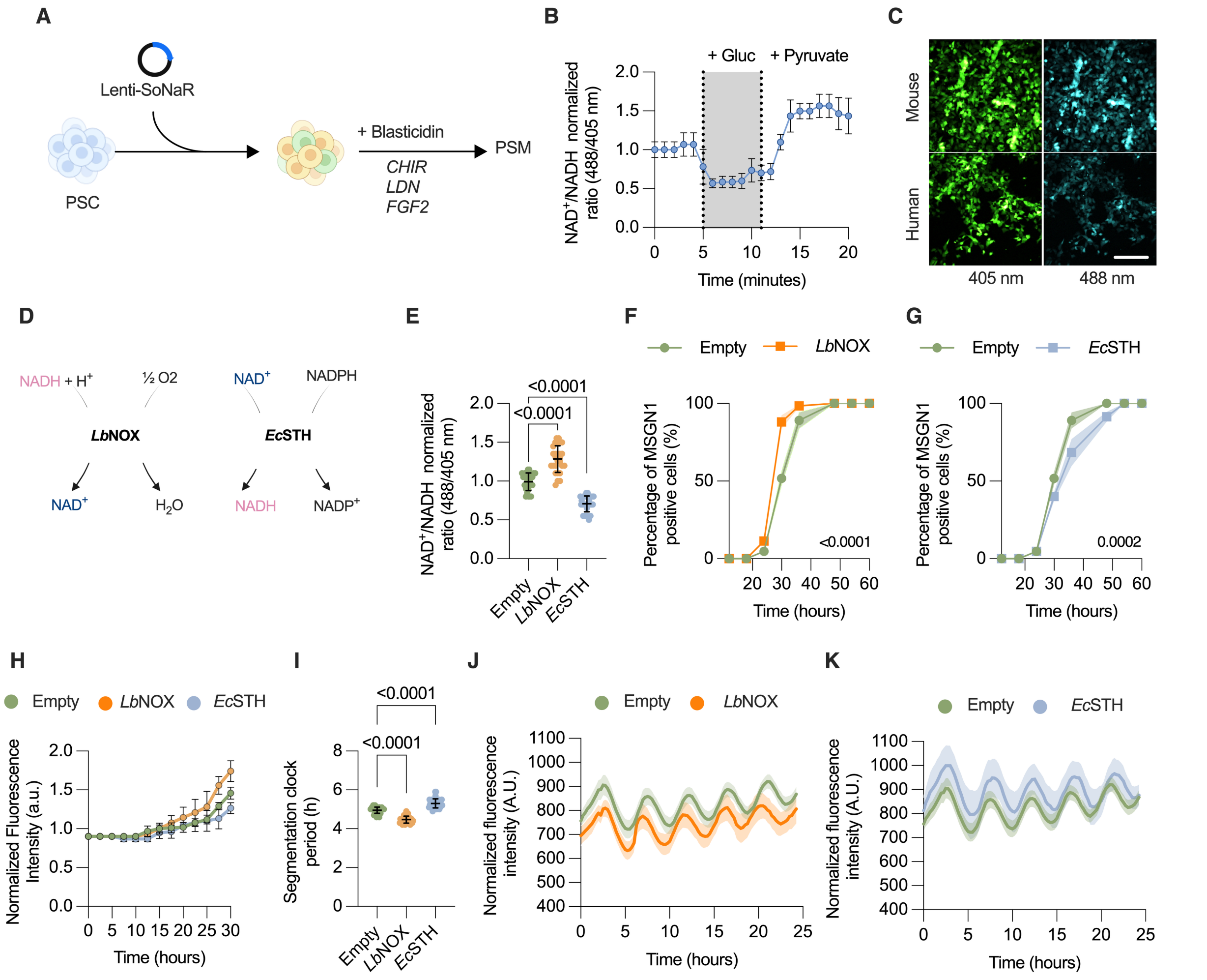

**Fig. S3 Validation of NAD$^+$/NADH redox sensor and comparative analysis of NAD$^+$ metabolism in mouse and human PSM.** (**A**) Experimental workflow for SoNar-based redox imaging. Pluripotent stem cells (PSCs) were transduced with Lenti-SoNar, selected with blasticidin, and differentiated to PSM using CHIR99021, LDN193189, and FGF2. (**B**) Dynamic validation of the SoNar sensor in human control PSM cells showing reversible changes in the NAD$^+$/NADH ratio upon sequential addition of glucose and pyruvate (n = 3 independent experiments). Data are presented as mean ± SD. (**C**) Representative 405- and 488-nm excitation images of mouse and human PSM cells expressing SoNar, showing a higher NAD$^+$/NADH ratio (more oxidized redox state) in mouse PSM. Scale bar, 50 µm. (**D**) Schematic of the redox-modulating enzymes *Lb*NOX and *Ec*STH, which drive NAD(H) oxidation or reduction, respectively. (**E**) Expression of *Lb*NOX or *Ec*STH oppositely shifts the intracellular NAD$^+$/NADH balance in human pluripotent stem cell-derived PSM cells as measured by ratiometric imaging of SoNar excitation (488/405nm), validating their functional activity (n > 3 independent replicates; > 200 cells per replicate; mean ± s.d.; one-way ANOVA with Dunnett's post hoc test, $P < 0.0001$). (**F–G**) Quantification of the percentage of MSGN1-Venus$^+$ cells over time shows that *Lb*NOX accelerates and *Ec*STH delays PSM fate acquisition relative to empty-vector controls in differentiating human iPSCs, linking redox state to developmental tempo (n = 3 independent replicates; repeated-measures two-way ANOVA with Tukey correction; i, $P < 0.0001$; j, $P = 0.0002$). (**H**) Changes in fluorescence intensity of the MSGN1 reporter over time in differentiating human cells expressing empty vector, *Lb*NOX, or *Ec*STH (n = 3 independent experiments; mean ± SD). (**I**) Segmentation clock period in human pluripotent stem cell-derived PSM cells upon redox perturbation: the period is shortened by NAD(H) oxidation (*Lb*NOX expression) and prolonged by NAD(H) reduction (*Ec*STH expression) compared to an empty vector control (n > 3 independent replicates; > 30 cells per replicate; mean ± s.d.; one-way ANOVA with Dunnett's post hoc test, $P < 0.0001$). (**J–K**) Representative oscillatory HES7 fluorescence intensity traces in human PSM cells expressing empty vector, *Lb*NOX (j), or *Ec*STH (k) (n = 3 independent experiments; mean ± SD).

**Fig. S4.**

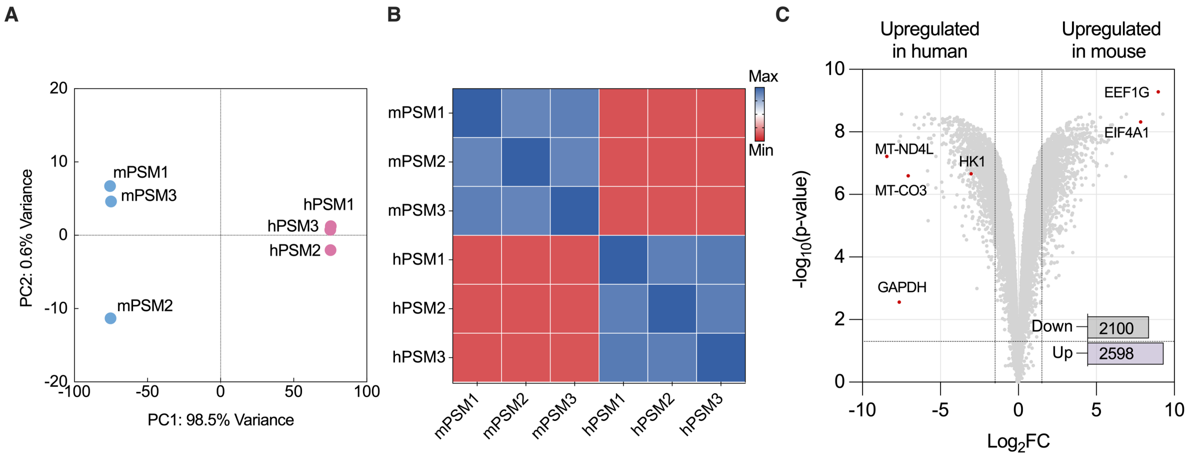

**Fig. S4 Comparative transcriptomic analysis between mouse and human PSM cells.** (**A**) Principal component analysis (PCA) of bulk RNA-seq samples showing clear species segregation along PC1 (98.5 % variance) between mouse and human PSM cells derived from pluripotent stem cells. (**B**) Sample-to-sample distance heatmap illustrating within-species clustering and high reproducibility across biological replicates in bulk RNA-sequencing data of mouse and human PSM cells derived from pluripotent stem cells. (**C**) Volcano plot of differentially expressed genes between mouse and human PSM cells *in vitro*, highlighting up-regulation of oxidative phosphorylation components (*MT-ND4L*, *MT-CO3*) and glycolytic enzyme *HK1* in human, and translation-related genes (*EEF1G*, *EIF4A1*) in mouse. Numbers indicate total up- and down-regulated genes.

**Fig. S5**

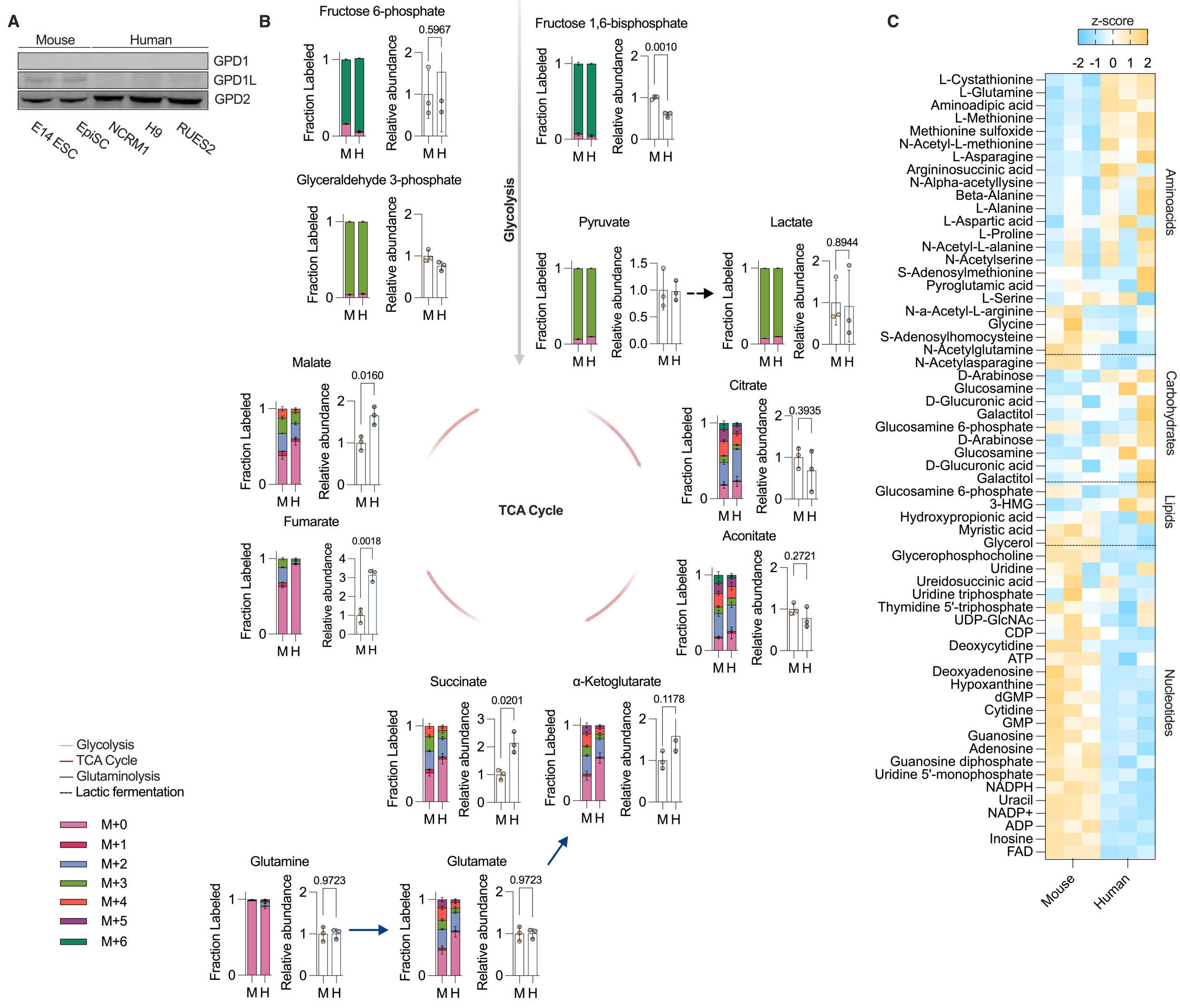

**Fig. S5 Comparative metabolomic profiling between mouse and human PSM.** (**A**) Immunoblotting of GPD1, GPD1L, and GPD2 in PSM cells derived from naïve E14 mouse embryonic stem cells (E14 ESC), mouse epiblast stem cells (EpiSC), human induced pluripotent stem cells (NCRM1), and human embryonic stem cell lines (H9 and RUES2). (**B**) Steady-state abundance (right) and [U-$^{13}C_6$]glucose labeling (left) of key glycolytic and TCA-cycle intermediates in mouse (M) and human (H) *in vitro*-derived PSM cells. Mouse cells show higher fractional labeling and relative abundance of several glycolytic (e.g., fructose-1,6-bisphosphate) and TCA-cycle intermediates (malate, fumarate, succinate), indicating increased carbon flux through oxidative metabolism (n = 3 independent experiments per species; mean ± SD; left, stacked bar plots show isotopologue distribution; right, bar plots show relative metabolite abundance; unpaired two-tailed t-test). (**C**) Heat map summarizing relative abundance (z-score) of amino acids, carbohydrates, lipids, and nucleotides in mouse and human PSM cells *in vitro,* determined by LC–MS (n = 3 independent experiments per species). Mouse cells exhibit elevated levels of TCA-linked and $NAD^+$-associated metabolites, consistent with enhanced oxidative and biosynthetic capacity.

**Fig. S6.**

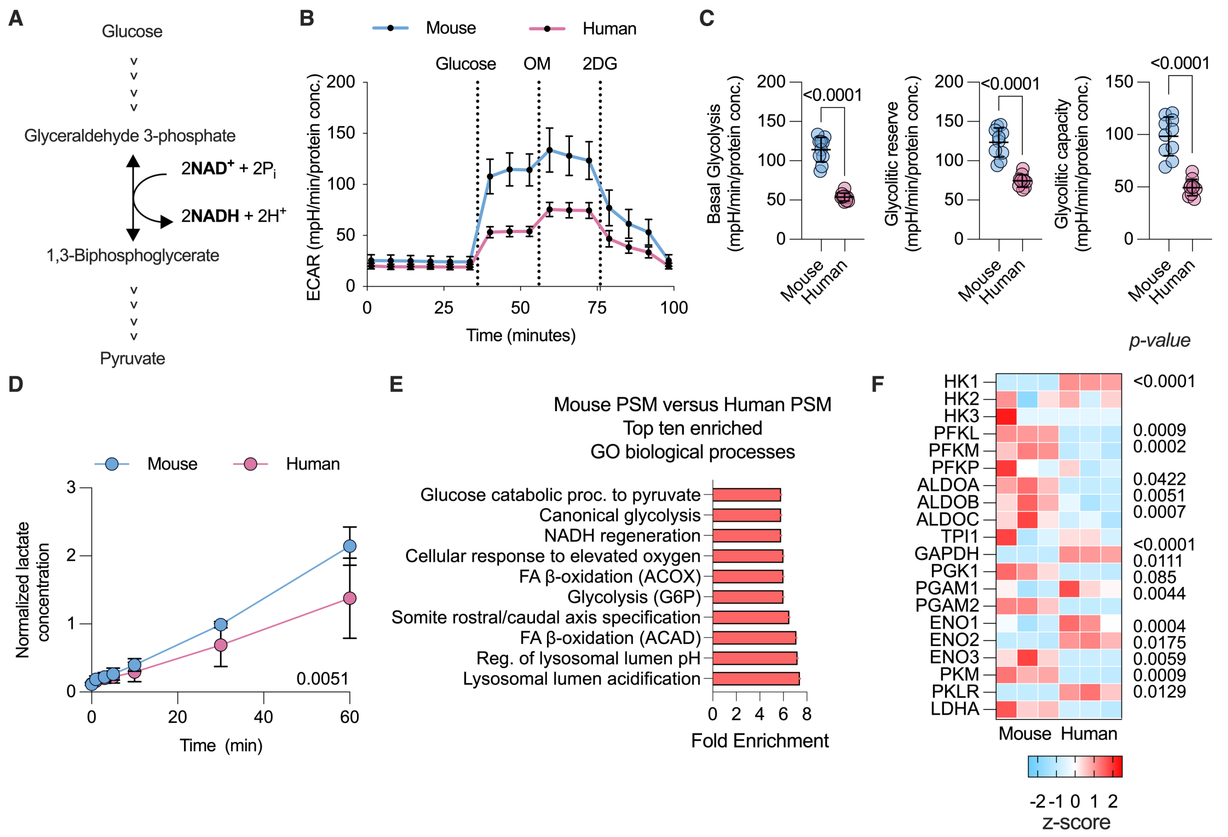

**Fig. S6. Reduced glycolytic throughput in human PSM cells.** (**A**) Schematic of the glycolytic reaction catalyzed by glyceraldehyde-3-phosphate dehydrogenase, illustrating the generation of NADH from $NAD^+$ during the oxidation of glyceraldehyde-3-phosphate to 1,3-bisphosphoglycerate. (**B**) Extracellular acidification rate (ECAR) measurements in mouse and human PSM cultures *in vitro* following sequential addition of glucose, oligomycin (OM), and 2-deoxy-D-glucose (2DG), showing higher glycolytic flux and capacity in mouse PSM (n = 8 biologically independent samples per species; data are presented as mean ± SD).
(**C**) Extracellular flux measurements of basal glycolysis, glycolytic reserve, and glycolytic capacity in mouse and human pluripotent stem cell-derived PSM cells (n > 8 independent samples per species; mean ± s.d.; unpaired two-tailed t-test, $P < 0.0001$). (**D**) Time-course of lactate accumulation in culture supernatants normalized to baseline levels for mouse and human pluripotent stem cell-derived PSM cells, showing higher glycolytic output in mouse relative to human PSM (n = 3 independent replicates per species; mean ± SD; repeated-measures two-way ANOVA with Tukey correction, P = 0.0051). (**E**) Gene Ontology (GO) analysis of transcripts enriched in mouse relative to human PSM cells highlights pathways associated with glycolysis, NADH regeneration, and β-oxidation (statistical analysis detailed in Methods). (**F**) Heat map showing expression (z-score) of glycolytic genes across species, revealing coordinated up-

regulation of key glycolytic enzymes in mouse cells (n = 3 independent replicates).

**Fig. S7.**

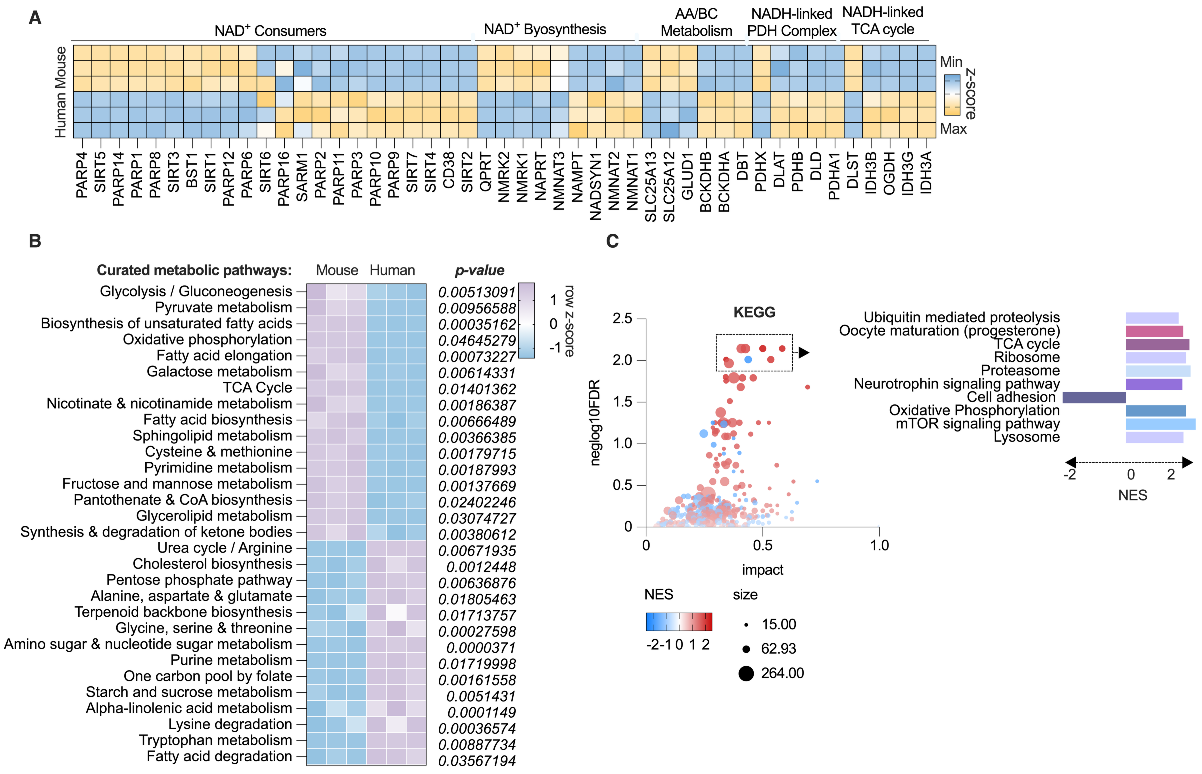

**Fig. S7 Transcriptomic comparison of mouse and human presomitic mesoderm (PSM) reveals metabolic pathway divergence.** (**A**) Heat map based on bulk RNA-sequencing data of transcripts encoding enzymes involved in NADH-linked reactions, $NAD^+$ biosynthesis, and $NAD^+$ consumption pathways in mouse and human PSM derived *in vitro* (n = 3 independent replicates per species). (**B**) Gene-set enrichment analysis (GSEA) of curated metabolic pathways showing enrichment of glycolysis/gluconeogenesis, pyruvate metabolism, and nicotinamide metabolism among top differentially regulated pathways between mouse and human pluripotent stem cell-derived PSM cells. (**C**) KEGG pathway enrichment analysis visualized by bubble plot (left) and normalized enrichment score (NES) summary (right), showing differential regulation of oxidative phosphorylation, TCA cycle, and mTOR-related pathways between mouse and human pluripotent stem cell-derived PSM cells.

**Fig. S8.**

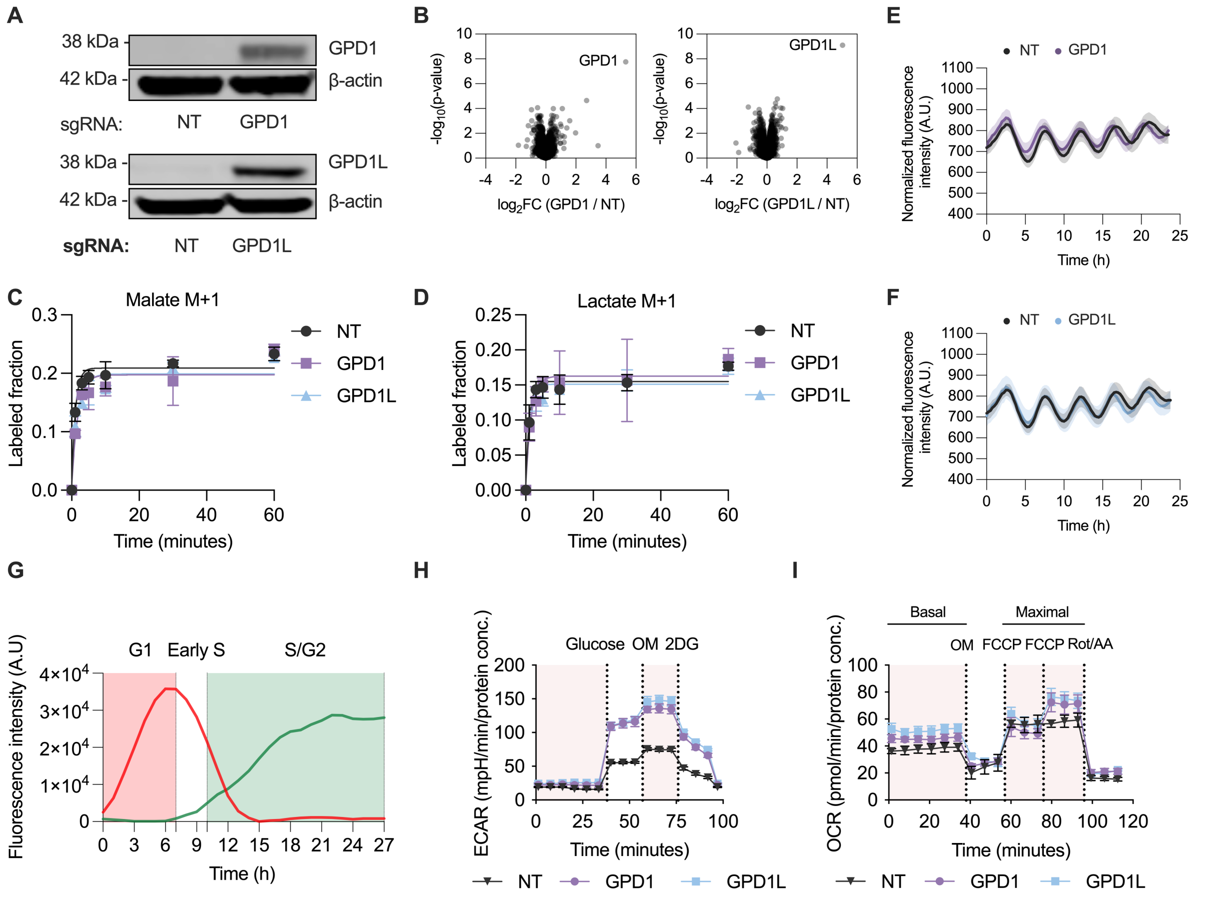

**Fig. S8 Metabolic consequences of GPD1 and GPD1L activation.** (**A**) Immunoblot validation of GPD1 and GPD1L overexpression in human PSM cells; β-actin served as loading control. (**B**) Volcano plots based on TMT-proteomics showing upregulation of GPD1 (left) and GPD1L (right) following targeted overexpression. (**C–D**) Time-course of deuterium enrichment (M+1) in malate (**C**) and lactate (**D**) following $[4\text{-}^2H]$glucose labeling in human PSM cells, showing that GPD1 or GPD1L overexpression does not alter labeling kinetics of these NADH-dependent metabolites compared to non-targeting (NT) controls (n = 3 independent experiments per condition; mean ± SD; repeated-measures two-way ANOVA, not significant for both metabolites). (**E–F**) Representative oscillatory traces of HES7–Achilles reporter activity in GPD1 (e) and GPD1L (f) overexpressing human PSM cells compared to non-targeting (NT) controls (n = 3 independent replicates; mean ± SEM). (**G**) Representative trace of a single FUCCI-reporter cell showing dynamic fluorescence corresponding to cell-cycle progression (red = G1 phase, green = S/G2 phase). (**H**) Extracellular acidification rate (ECAR) profiles measured by Seahorse flux analysis in control (NT) and GPD1/GPD1L-overexpressing human PSM cells, showing enhanced glycolytic activity and capacity upon GPD1 or GPD1L overexpression. Glucose, oligomycin (OM) and 2-deoxy-D-glucose (2DG) were sequentially injected (n = 8 biologically independent samples per condition; mean ± SD). (**I**) Oxygen consumption rate (OCR) measurements in control (NT) and GPD1/GPD1L-overexpressing human PSM cells, indicating increased basal and maximal respiration relative to NT controls. Oligomycin (OM), carbonyl cyanide p-trifluoromethoxyphenylhydrazone (FCCP) and rotenone plus antimycin A (Rot and AA) were sequentially injected (n = 8 biologically independent samples per condition; mean ± SD).

**Fig. S9.**

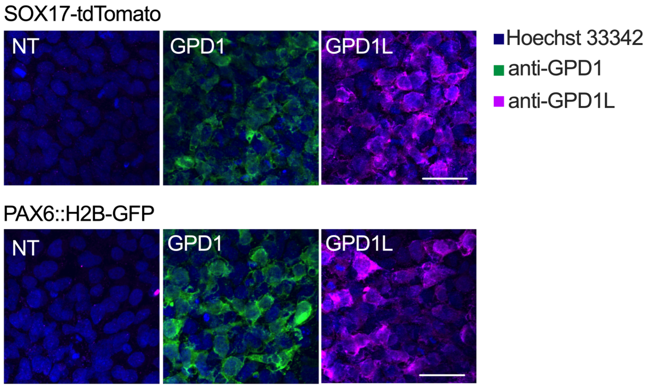

**Fig. S9. Representative images of DE and NPC cells overexpressing GPD1 or GPD1L.** Representative immunofluorescence images of definitive endoderm (DE; SOX17-tdTomato reporter, top) and neural progenitor cells (NPC; PAX6-H2B-GFP reporter, bottom) transduced with GPD1- or GPD1L-sgRNAs. Nuclei are labeled with Hoechst 33342 (blue). Overexpression was confirmed by staining with anti-GPD1 (green) or anti-GPD1L (magenta). NT, non-targeting control. Scale bar, 100 µm.

### Table S1. Cell lines

| Species | Cell line | Reporter/transgene | Integration method | Source |
|---|---|---|---|---|
| **Human** | *NCRM1* Induced Pluripotent Stem Cells | N/A | N/A | NINDS Stem Cell and Data Repository |
| | | *HES7-Achilles* | CRISPR | Olivier Pourquie, Brigham and Women's Hospital |
| | | *HES7-Achilles; CAG-dCas9-VP64* | CRISPR | This study |
| | | *CAG-FUCCI* | CRISPR | This study |
| | | *CAG-FUCCI; CAG-dCas9-VP64* | Lentivirus | This study |
| | | *EF1a-cytoSoNAR* | Lentivirus | This study |
| | | *CAG-dCas9-VP64; EF1a-cytoSoNar* | Lentivirus | This study |
| | *hiPS-11a* Induced Pluripotent Stem Cells | N/A | N/A | Harvard Stem Cell Institute |
| | | *MSGN1-Venus* | CRISPR | Olivier Pourquie, Brigham and Women's Hospital |
| | | *MSGN1-Venus; CAG-dCas9-VP64* | CRISPR | This study |
| | *H9* Embryonic Stem Cells | N/A | N/A | WiCell |
| | | *PAX6-GFP* | CRISPR | Lorenz Studer, Memorial Sloan Kettering Cancer Center |
| | | *PAX6-GFP; CAG-dCas9-VP64* | CRISPR | This study |
| | *RUES2* Embryonic Stem Cells | *Germ Layer Reporter (GLR): SOX17-tdTomato; Bra-mCerulean; SOX2-mCitrine* | CRISPR | Ali Brivanlou, Rockefeller University |
| | | *Germ Layer Reporter (GLR): SOX17-tdTomato; Bra-mCerulean; SOX2-mCitrine; CAG-dCas9-VP64* | Lentivirus | This study |
| | HEK *293T* | N/A | N/A | Takara, cat. no. 632180 |
| **Mouse** | *E14* Embryonic Stem Cells | pMsgn1-Venus | PiggyBac | Olivier Pourquie, Brigham and Women's Hospital |
| | Epiblast Stem Cells | N/A | N/A | Jun Wu, University of Texas Southwestern Medical Center |
| | | *EF1a-cytoSoNar* | Lentivirus | This study |
| | | *Hes7-Achilles* | CRISPR | This study |
| | | *Msgn1-Achilles* | CRISPR | This study |

### Table S2. Single guide RNAs

| Application | Target | Sequence |
|---|---|---|
| CRISPRa | *hGPD1* | GCCAGACTCTCTATCTCCCT |
| CRISPRa | *hGPD1L* | AGGCGTGCGCAGTGGTTCTT |
| CRISPRa | *hNT* | GTATTACTGATATTGGTGGG |
| Knock-in | *mMsgn1* | GGCACAACTCACACACTCTG |
| Knock-in | *mHes7* | GTCTCCAAAACGCGGGCGGT |
| Knock-in | *AAVS1* locus | CCGGTGTTGGAAGGATGAGGAAAT |

### Table S3. List of primers.

| Primer | Sequence | Source |
|---|---|---|
| **Amplify_sgRNA_fwd** | AGGCACTTGCTCGTACGACG | Sanson et al. 2018(*60*) |
| **Amplify_sgRNA_rev** | ATGTGGGCCCGGCACCTTAA | Sanson et al. 2018(*60*) |
| **mHes7_out_fwd** | CGCCTGATCAATGGGCTTCAG | Diaz-Cuadros et al. 2020(*4*) |
| **mHes7_out_rev** | TGGAGAGCAGGCCTAAAGGTG | Diaz-Cuadros et al. 2020(*4*) |
| **Achilles_in_fwd** | TCACTCTCGGCATGGACGAG | Diaz-Cuadros et al. 2020(*4*) |
| **Achilles_in_rev** | CGCTGAACTTGTGGCCGTTTAC | Diaz-Cuadros et al. 2020(*4*) |
| **mMsgn1_out_fwd** | GAATCAAAGCTCAGGCGACTCAC | This study |
| **mMsgn1_out_rev** | AGGTACCCAGCAGGACAGTG | This study |
| **dna_803 (AAVS1)** | TCGACTTCCCCTCTTCCGATG | Oceguera-Yanez et al. 2016(*76*) |
| **dna_804 (AAVS1)** | GAGCCTAGGGCCGGGATTCTC | Oceguera-Yanez et al. 2016(*76*) |
| **dna_183 (AAVS1)** | CTCAGGTTCTGGGAGAGGGTAG | Oceguera-Yanez et al. 2016(*76*) |

**Table S4. List of antibodies**

| Antibody | Source | Cat number | Dilution |
|---|---|---|---|
| **GPD1 polyclonal antibody** | Proteintech | 27943-1-AP | 1:1000 (immunoblotting), 1:200 (immunofluorescence) |
| **GPD1L polyclonal antibody** | Proteintech | 17263-1-AP | 1:1000 (immunoblotting), 1:200 (immunofluorescence) |
| **GPD2 monoclonal antibody** | Proteintech | 68174-1-Ig | 1:1000 (immunoblotting) |
| **BETA-ACTIN (8h10d10) mouse monoclonal antibody #3700** | Cell Signaling | # 3700 | 1:10000 (immunoblotting) |
| **NANOG monoclonal antibody** | Proteintech | 67255-1-Ig | 1:200 (immunofluorescence) |
| **Anti-SOX2 antibody** | Millipore Sigma | AB5603 | 1:200 (immunofluorescence) |
| **OCT3/4** | Santa Cruz | sc-5279 | 1:500 (immunofluorescence) |
| **Anti-TBX6 antibody** | Abcam | ab38883 | 1:200 (immunofluorescence) |
| **TBXT** | R&D | AF2085 | 1:200 (immunofluorescence) |
| **SOX17** | Proteintech | 24903-1-AP | 1:200 (immunofluorescence) |
| **ANTI-PAX-6 ANTIBODY** | BioLegend | 901302 | 1:200 (immunofluorescence) |
